# Molecular interplay between HURP and Kif18A in mitotic spindle regulation

**DOI:** 10.1101/2024.04.11.589088

**Authors:** Juan M. Perez-Bertoldi, Yuanchang Zhao, Akanksha Thawani, Ahmet Yildiz, Eva Nogales

## Abstract

During mitosis, microtubule dynamics are regulated to ensure proper alignment and segregation of chromosomes. The dynamics of kinetochore-attached microtubules are regulated by hepatoma-upregulated protein (HURP) and the mitotic kinesin-8 Kif18A, but the underlying mechanism remains elusive. Using single-molecule imaging *in vitro*, we demonstrate that Kif18A motility is regulated by HURP. While sparse decoration of HURP activates the motor, higher concentrations hinder processive motility. To shed light on this behavior, we determined the binding mode of HURP to microtubules using Cryo-EM. The structure reveals that one HURP motif spans laterally across β-tubulin, while a second motif binds between adjacent protofilaments. HURP partially overlaps with the microtubule-binding site of the Kif18A motor domain, indicating that excess HURP inhibits Kif18A motility by steric exclusion. We also observed that HURP and Kif18A function together to suppress dynamics of the microtubule plus-end, providing a mechanistic basis for how they collectively serve in spindle length control.

## Introduction

Proper segregation of genetic material during cell division relies on the organization of microtubule filaments into a bipolar spindle. Polarity and polymerization dynamics of spindle microtubules are regulated by a plethora of microtubule-associated proteins (MAPs) and molecular motors to form specialized sub-structures within the spindle. Kinetochore-fibers (K-fibers), made of parallel arrays of kinetochore-bound (K-microtubules), and non-kinetochore-bound microtubules, are an example of spindle specialization. During metaphase, the chromosomes attach to K-fibers and experience oscillatory movements to facilitate the alignment of chromosomes at the metaphase plate and generate the pulling forces on the microtubules that enable chromosome segregation^1–4^. K-microtubules remain tightly bound to the kinetochores during the growth and shrinking phases of the dynamic plus-ends and the spindle globally maintains a constant steady-state length during metaphase^5–7^. Because improper chromosome alignment can lead to aneuploidy, cancer, and birth defects^8–11^, it is important to understand the mechanisms regulating the properties and function of K-fibers. Yet, the molecular mechanism of how microtubule properties are altered to robustly engage K-fibers with kinetochores throughout cell division is not well understood.

K-fibers recruit specific MAPs and motors to promote microtubule bundling and modulate their plus-end dynamics. Hepatoma upregulated protein (HURP) is a spindle assembly factor (SAF) that localizes to the chromatin-proximal region of K-fibers in a process mediated by Ran-GTP signaling^12–14^. Through its stabilizing and bundling activities, HURP regulates K-fiber dynamics and spindle morphology, contributing to chromosomal movements and alignment^15,16^. Both depletion and overexpression of HURP lead to defective spindles that cannot align chromosomes effectively^14,17^. However, how HURP binds to microtubules and stabilizes their ends remains unexplored.

K-fibers also recruit Kif18A, a member of the kinesin-8 family that walks towards and accumulates at the plus-ends of K-microtubules^18–21^. Kif18A modulates directional switching of chromosome oscillations and the relative motion of sister kinetochores, termed breathing, by suppressing the plus-end growth of K-microtubules in a length-dependent manner^22–24^. The mechanism by which Kif18A regulates microtubule dynamics remains controversial^25^. An earlier study showed that Kif18A actively depolymerizes GMPCPP-stabilized microtubules *in vitro*^20^, whereas another study argued that Kif18A primarily controls microtubule length by acting as a capping protein and restraining growth rate, without necessarily inducing depolymerization^26^. Although Kif18A and its homologs have been studied *in vivo* and *in vitro*, the molecular understanding of how the motor accumulates at the plus-end of K-microtubules and modulates microtubule dynamics remains to be demonstrated. Interestingly, Kif18A-depleted cells exhibit similar phenotypes to those of HURP-depleted cells^17,21,27^, suggesting a relationship between the functions of these two proteins.

Studies in live cells suggested that the N-terminus of HURP binds to microtubules^28,29^ and regulates Kif18A localization on these tubulin polymers^17^. Here, we use total internal reflection fluorescence (TIRF) microscopy to reveal how Kif18A motility on microtubules is influenced by HURP *in vitro* and to show that these two proteins work together to modulate plus-end dynamics of microtubules. Kif18A is recruited to the microtubule lattice and activated by HURP, resulting in enhanced Kif18A motility. Yet, at high HURP concentrations, we observe antagonism between HURP and Kif18A. To better understand this mutual antagonism, we employed cryogenic electron microscopy (cryo-EM) to visualize how HURP and Kif18A bind to the microtubules at near-atomic resolution. Our structural studies reveal an overlap of the binding surfaces of the two proteins on the microtubule, resulting in the inhibition of KIF18A by excess HURP on the microtubule. We also show that the interplay between HURP and Kif18A at the plus-end modulates microtubule dynamics. This could provide a mechanism for K-microtubule stabilization and length control, ultimately impacting on chromosome congression within the mitotic spindle.

## Results

### HURP regulates Kif18A motility in a concentration-dependent manner

To explore how HURP regulates Kif18A motility, we recombinantly expressed full-length and truncated HURP constructs C-terminally tagged with an enhanced green fluorescent protein (HURP^1–173^-eGFP, HURP^1-285^-eGFP, HURP^1-400^-eGFP and HURP^1-846^-eGFP (full-length HURP)) (Figure 1 A). We used TIRF microscopy to quantify microtubule binding of HURP (Figure 1 B). All four HURP constructs bound to microtubules (Figure 1 C). The dissociation constant (K_d_) of HURP^1-285^ binding to taxol-stabilized microtubules was ∼0.6 µM at physiological salt concentration (150 mM) (Figure 1 D).

**Figure 1.**
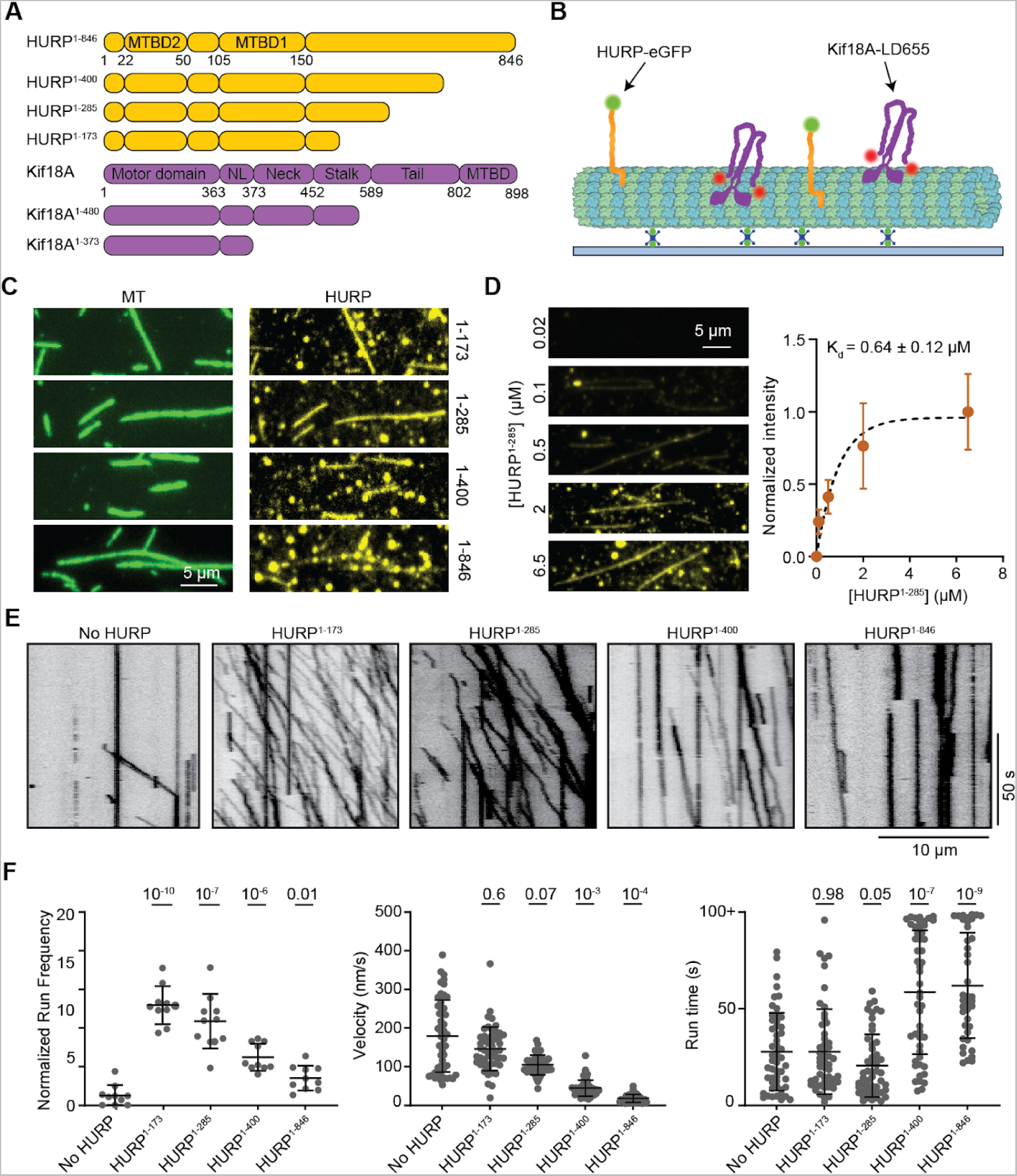
HURP contains different elements that can activate or decelerate Kif18A motility. **A**. Domain organization of full-length HURP and Kif18A, and different HURP and Kif18A truncations used in this study (NL: neck-linker). **B.** Schematic of the *in vitro* reconstitution of Kif18A motility on surface-immobilized microtubules in the presence of HURP. **C.** Representative fluorescence images showing microtubule binding of HURP constructs. **D.** (Left) Representative images showing HURP^1-285^ binding to microtubules at different concentrations. (Right) Quantification of HURP^1-285^ binding to microtubules. The center circle and whiskers represent the mean and S.D., respectively. Kd is determined from a fit to binding isotherm (dashed curve, from left to right, n = 49, 50, 50, 24, 52 microtubules). **E.** Representative kymographs showing motility of Kif18A in the presence of different HURP constructs at 1-2 µM. **F.** Normalized run frequency (n = 10 kymographs for each condition), velocity and run time (from left to right, n = 52, 52, 52, 50, 40 motors) of Kif18A in the presence of different HURP constructs. The center line and whiskers represent the mean and S.D., respectively. P values were calculated from a two-tailed t-test, compared to the no HURP condition.

We next assayed the motility of human Kif18A on surface-immobilized microtubules in the presence and absence of near saturating (1-2 µM) HURP concentrations (Figure 1 B). HURP^1-173^ and HURP^1-285^ substantially enhanced the run frequency of Kif18A without significantly altering its velocity or run time (Figure 1 E-F), suggesting that HURP contains a Kif18A-activating site between amino acids 1-173. The stimulatory effect of HURP on Kif18A motility appeared specific, as HURP^1-285^ did not activate kinesin-1 Kif5B (Supplementary Figure 1), and Kif18A activation was not observed with the MAPs doublecortin, MAP7, or tau (Supplementary Figure 2). HURP^1-400^ and HURP^1-846^ also increased the landing of Kif18A on the microtubule. However, the effect of HURP^1-400^ and HURP^1-846^ on the run frequency of Kif18A was not as pronounced as with the shorter HURP constructs because HURP^1-400^ and HURP^1-846^ reduced Kif18A velocity several fold, resulting in more motors being counted as stationary (Figure 1 E-F).

We then investigated how HURP^1-846^ affects Kif18A motility at lower concentrations (0 - 0.5 µM). Kif18A run frequency exhibited a biphasic behavior with an initial activation phase followed by a decrease in the run frequency at near saturating concentrations of HURP^1-846^. In comparison, Kif18A velocity displayed a steady decrease and the run time steadily increased under increasing concentrations of HURP^1-846^ (Figure 2 A).

**Figure 2.**
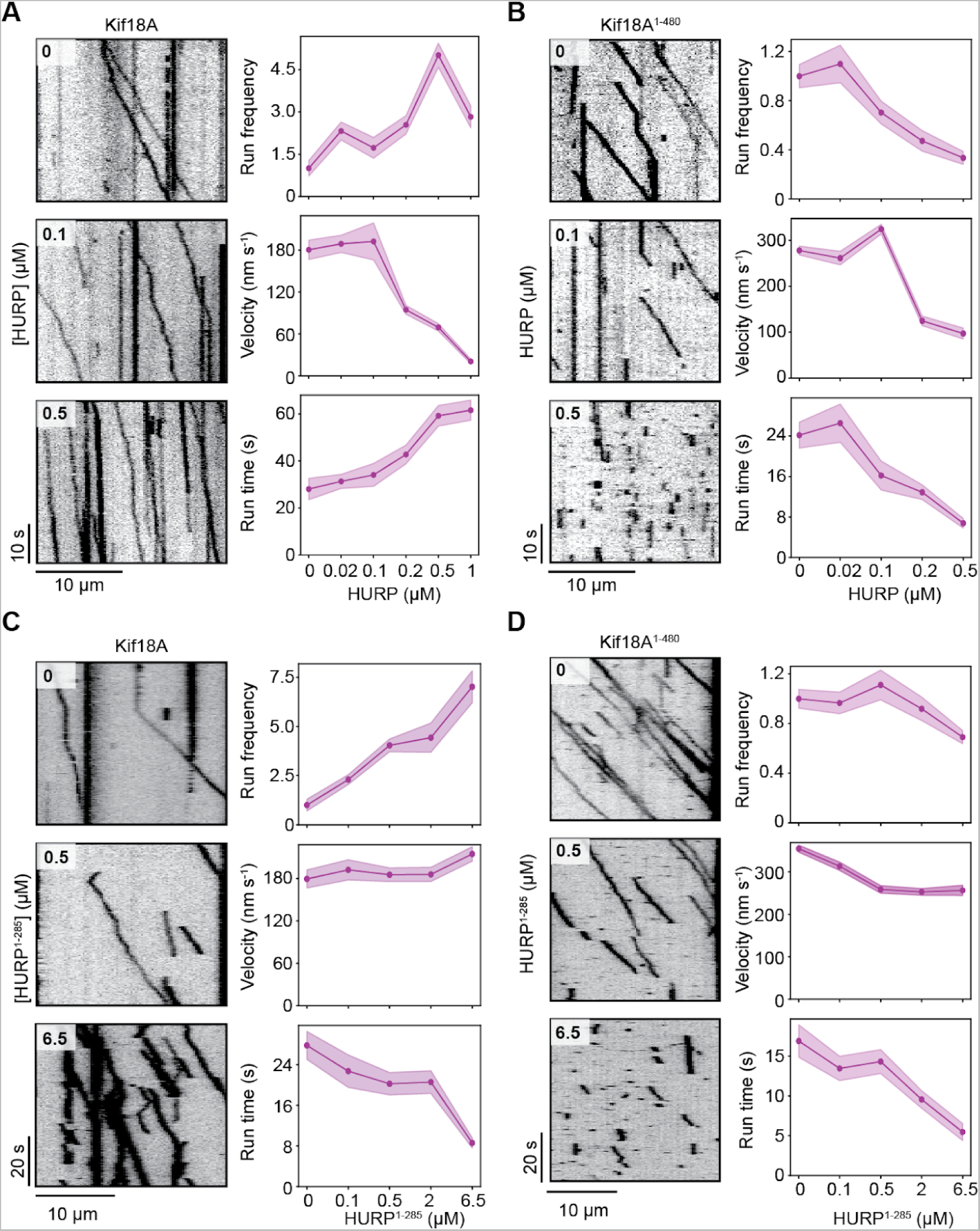
HURP differentially affects Kif18A motility in a concentration-dependent manner. **A**. (Left) Representative kymographs showing the motility of full-length Kif18A in the presence of 0, 0.1, and 0.5 µM HURP^1-846^. (Right) Normalized run frequency (n = 10 kymographs for each condition), velocity and run time (from left to right, n = 32, 50, 33, 48, 50 motors) of Kif18A for different HURP concentrations. **B**. (Left) Representative kymographs showing the motility of Kif18A^1-480^ in the presence of 0, 0.1, and 0.5 µM HURP^1-846^. (Right) Normalized run frequency (n = 10 kymographs for each condition), velocity and run time (n = 25 motors for each condition) of Kif18A^1-480^ for different HURP concentrations. **C**. (Left) Representative kymographs showing the motility of full-length Kif18A in the presence of 0, 0.5, and 6.5 µM HURP^1-285^. (Right) Normalized run frequency (n = 10 kymographs for each condition), velocity and run time (from left to right, n = 52, 44, 54, 52, 52 motors) of Kif18A for different HURP^1-285^ concentrations. **D**. (Left) Representative kymographs showing the motility of Kif18A^1-480^ in the presence of 0, 0.5, and 6.5 µM HURP^1-285^. (Right) Normalized run frequency (n = 10 kymographs for each condition), velocity and run time (from left to right, n = 52, 51, 52, 52, 52 motors) of Kif18A^1-480^ for different HURP^1-285^ concentrations. In **A-D** (right) the line and shadows represent the mean and S.E., respectively.

We next removed the C-terminal tail of Kif18A (Figure 1 A) to explore its possible role in the activation and HURP-mediated regulation of Kif18A motility. HURP^1-846^ also reduced the velocity of Kif18A^1-480^, suggesting that the slow down effect does not involve the C-terminal tail of Kif18A. Notably, Kif18A^1-480^ run frequency did not exhibit the activation phase observed with the full-length motor. Instead, its run frequency, velocity, run time and run length decreased as HURP^1-846^ concentration was increased (Figure 2 B, Supplementary Figure 3).

We also analyzed the motility of full-length Kif18A and Kif18A^1-480^ under 0-6.5 µM HURP^1-285^. The addition of this shorter HURP construct resulted in up to a seven-fold increase in Kif18A run frequency, showing that HURP^1-285^ enhances Kif18A motility in a concentration-dependent manner. On the other hand, HURP^1-285^ reduced the run time of Kif18A at higher concentrations (Figure 2 C), which is consistent with HURP^1-278^ overexpression mimicking a phenotype of Kif18A depletion *in vivo*^17^. Interestingly, in the absence of HURP, the tail-truncated Kif18A^1-480^ exhibited significantly more frequent runs than full-length Kif18A (Figure 2 A-D), suggesting that the motor may be partially autoinhibited by its tail, analogous to other kinesins^30–34^. HURP^1-285^ binding to microtubules did not significantly change the run frequency or velocity of Kif18A^1-480^, but substantially decreased the run time of the motor (Figure 2 D). These results are compatible with HURP interacting with the tail of full-length Kif18A, recruiting the motor to the microtubules, and rescuing it from autoinhibition to activate its motility.

Collectively, our functional studies suggest that the 285-400 segment of HURP decelerates Kif18A motility. Since the effect is observed for the motor even in the absence of its tail, it is possible that this HURP segment interacts with a region constrained to amino acids 1-480 of Kif18A. Unlike the 1-173 segment of HURP that appears to release auto-inhibition and activate motility, this interaction markedly reduces Kif18A’s velocity.

### HURP bridges tubulin subunits across protofilaments

To further understand the inhibition of Kif18A motility observed at higher HURP concentrations, we visualized how HURP interacts with the microtubule surface. A previous study identified two distinct microtubule binding domains (MTBDs) on the N-terminus of HURP: MTBD1 is the constitutive high-affinity interaction site (HURP^105-150^) whereas MTBD2 (HURP^22–50^) has weaker microtubule affinity and is regulated by importin-β^28^ (Figure 1 A). Our TIRF imaging assays revealed that HURP^1-285^ binds to taxol-stabilized microtubules with a K_d_ of ∼0.033 µM in the absence of added salt (Figure 3 A-B). We next used cryo-EM to determine the structure of HURP^1-285^ bound to taxol-stabilized microtubules under saturating conditions. After following a seam-determination procedure^35^ and exploiting the pseudo-symmetry of the microtubule to expand the effective number of particles, we generated a 3.1 Å cryo-EM density map (Figure 3 C, Supplementary Figure 4, Supplementary Figure 5 A) that allowed us to manually model HURP MTBD1 *de novo,* identifying residues 87-132 within the constitutive binding site. No density corresponding to the HURP MTBD2 was present in the map, indicating that the interaction between this region and microtubules likely involves flexible elements in either or both, HURP or tubulin.

**Figure 3.**
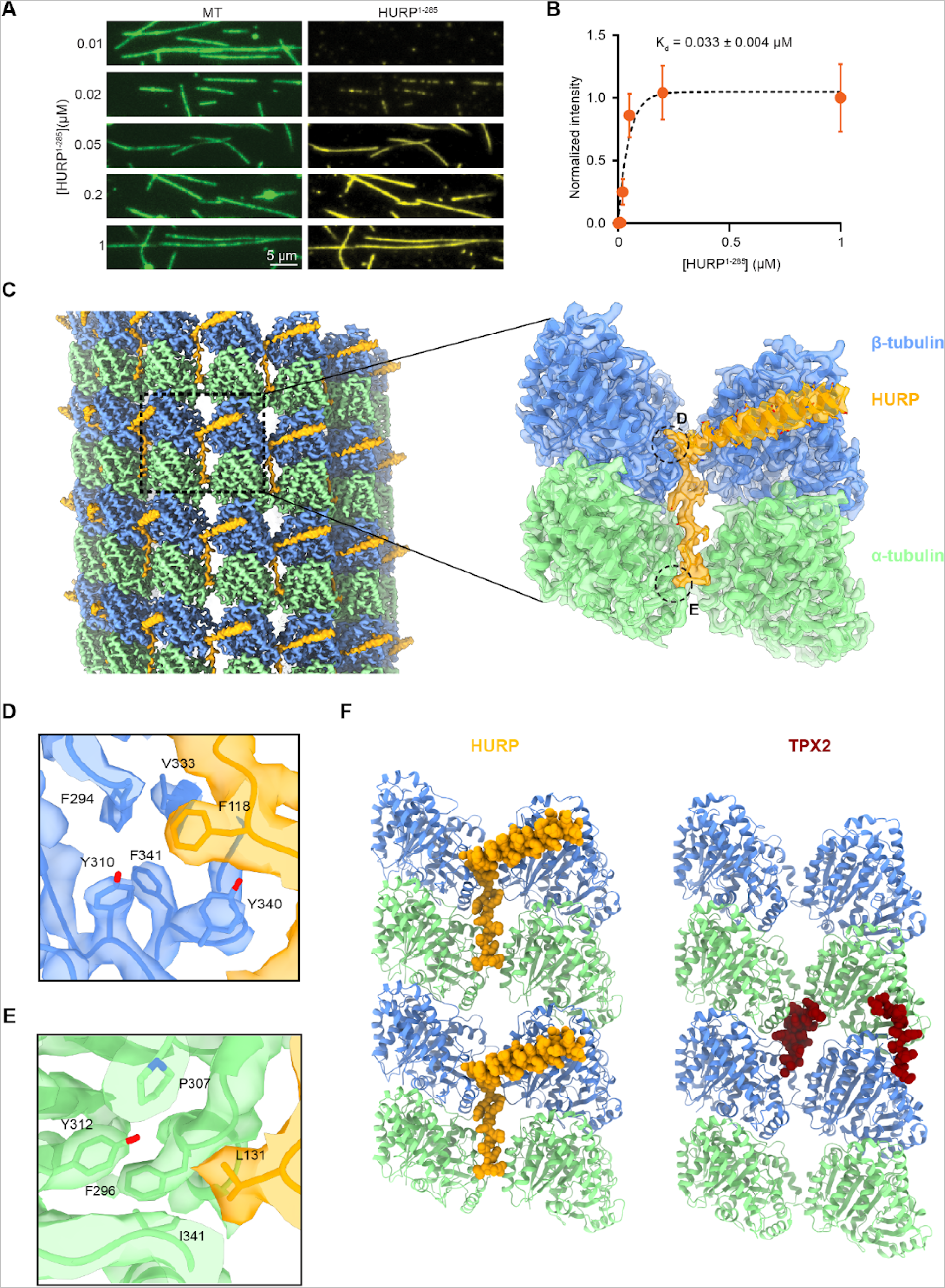
HURP interacts with microtubules through a dual binding site. **A.** Representative images showing microtubule decoration with different concentrations of HURP^1-285^ in the absence of added salt. **B.** Quantification of HURP^1-285^ binding to microtubules. The center line and whiskers represent the mean and S.D., respectively. The fit used to determine the Kd is determined from a fit to binding isotherm (dashed curve, n = 50 microtubules for each condition). **C.** (Left) Surface representation of the symmetrized microtubule-HURP cryo-EM density map generated with a mask around the entire microtubule. α-tubulin, 𝛽-tubulin and HURP are shown in green, blue and orange, respectively. The boxed region marks an area including two tubulin dimers and one HURP molecule. (Right) Final microtubule-HURP symmetry-expanded map generated with a mask focusing on two neighboring tubulin dimers and a single HURP molecule. A single HURP molecule is shown for clarity. The refined model is shown in ribbon representation and the map displayed with transparency. HURP side chains are displayed with atom representation (orange: C, red: O, blue: N, yellow: S) **D-E.** Details of the interactions between HURP and tubulin by the extended region across protofilaments. **F.** Comparison between HURP (orange, this study) and TPX2 (dark red, PDB 6BJC) microtubule-bound structures.

Two structural motifs were observed for MTBD1. An α-helical density (HURP^87-114^) spans laterally across β-tubulin and establishes contacts (via L94, Y97, K98 and K101) with residues on β-tubulin H12 (E410 and M406) through hydrogen bonding and hydrophobic interactions (Supplementary Figure 6 A-C). Additionally, an extended loop, including residues 115-132, inserts in the inter-protofilament groove and contacts two laterally adjacent tubulins, thus stapling the protofilaments together. These interactions are mainly driven by aromatic and hydrophobic residues that insert in hydrophobic pockets on the tubulin subunits (Figure 3 D-E, Supplementary Figure 6 A, D-E). Most of the HURP residues that participate in these interactions are highly conserved among species (Supplementary Figure 7). The deep insertion in the inter-protofilament groove and the bridging interactions between adjacent tubulin subunits are consistent with HURP’s role as a microtubule stabilizing factor^14,16^. The microtubule bound structure of HURP resembles that of another spindle assembly factor, TPX2, which also contains a dual binding motif that establishes lateral and longitudinal contacts to staple tubulins together^36^ (Figure 3 F). This similarity could point to a shared molecular mechanism for these two critical players in microtubule stabilization for the regulation of K-fiber dynamics (see Discussion).

### HURP and Kif18A cannot simultaneously occupy the same tubulin dimer

Superimposing our HURP-microtubule model with a previously reported Kif18A-microtubule structure (PDB 5OCU)^37^ reveals a potential steric clash between the α-helical segment of HURP and the L8/β5 tubulin binding motif of the Kif18A motor domain. Such overlap would be consistent with our observation that Kif18A cannot walk processively towards the plus-end at high HURP concentrations. To test whether HURP and Kif18A can co-occupy the microtubule lattice, we generated a monomeric Kif18A construct containing the motor domain and neck linker (Figure 1 A) fused to SNAP-tag (Kif18A^1-373^-SNAP). Then, we used TIRF imaging to determine the co-decoration of microtubules with HURP^1-285^-eGFP and Kif18A^1-373^-SNAP labeled with an LD655 dye (Figure 4 A). The two proteins coated the microtubules with similar surface densities when mixed at equal concentrations. Microtubule-binding of HURP^1-285^ and Kif18A^1-373^ was negatively correlated (Pearson’s R = −0.83, Figure 4 B), suggesting that HURP and Kif18A compete against each other for the available binding sites on the microtubule.

**Figure 4.**
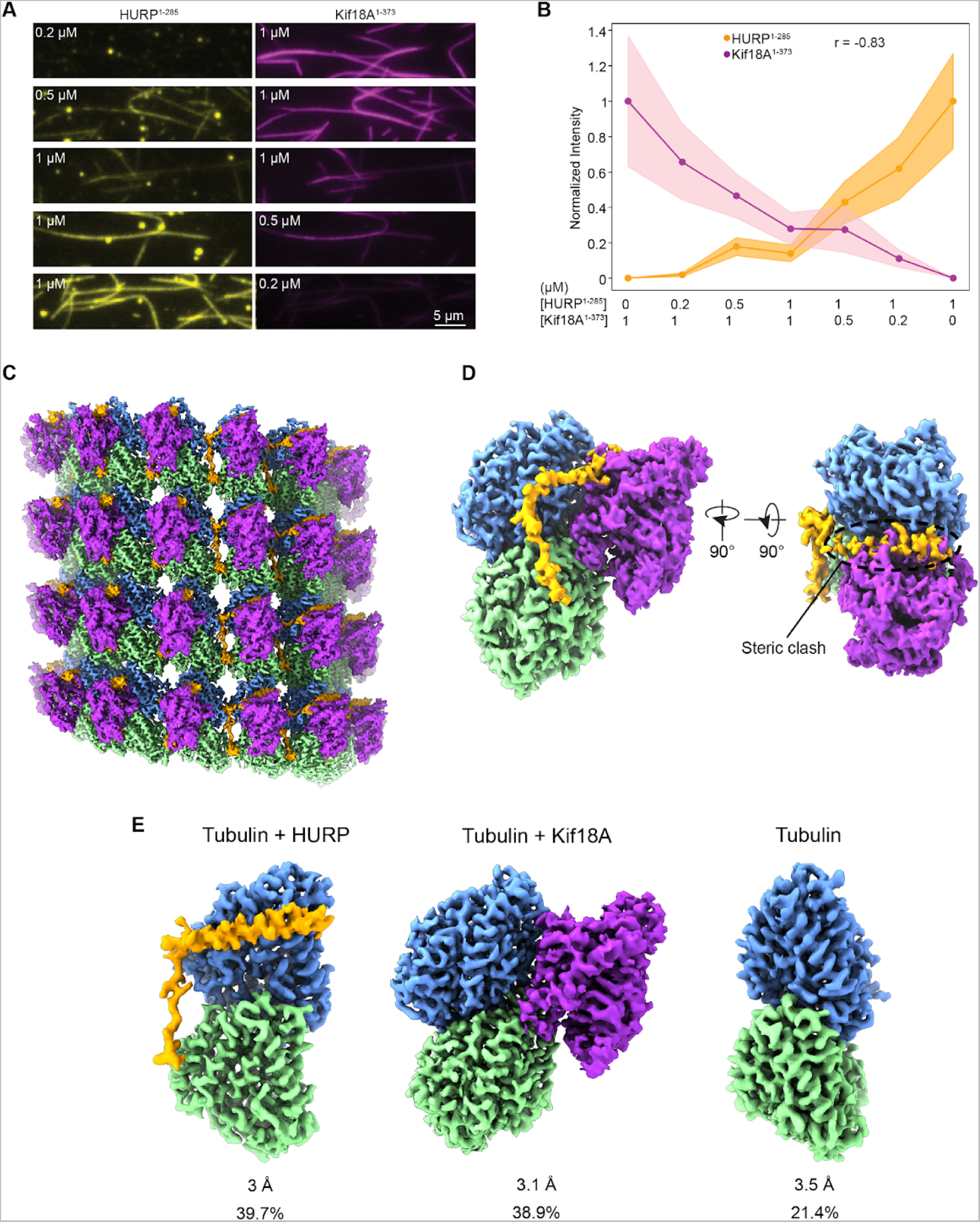
HURP binding site partially overlaps with Kif18A’s motor domain on the microtubules. **A.** Representative images of HURP^1-285^ -eGFP and Kif18A^1-373^-SNAP binding to microtubules for different ratios of the two proteins. **B.** Normalized fluorescence intensity for HURP^1-285^ and Kif18A^1-373^ for the protein ratios used in A. The center line and shadows represent the mean and S.E., respectively (n = 50 microtubules for each condition; r: Pearson’s correlation coefficient). **C.** Surface representation of the symmetrized microtubule-HURP-Kif18A^1-373^ cryo-EM density map generated with a mask around the entire microtubule. α-tubulin, 𝛽-tubulin, HURP and _mono_Kif18A are shown in green, blue, orange and purple, respectively. **D.** (Left) Final consensus reconstruction of the symmetry-expanded dataset generated with a mask focusing on two neighboring tubulin dimers (only one shown for clarity) and the bound HURP and Kif18A densities. (Right) Rotated volume showing the steric clash between HURP and Kif18A. The volumes are color coded as in C. **E.** Refined classes produced during alignment-free 3D classification shown in slightly different orientations for clarity. Resolution and particle class distribution for each are indicated.

To directly determine whether HURP and Kif18A exclude each other on microtubules, we determined the structure of microtubules co-decorated with HURP and Kif18A. To increase the likelihood of these proteins occupying the same or adjacent tubulins, we decorated taxol-stabilized microtubules with equimolar and near saturating concentrations of HURP and Kif18A. Processing of the cryo-EM images generated a consensus reconstruction for the co-decorated microtubule, showing clear density features for both HURP and Kif18A (Figure 4 C). The symmetry-expanded particle set was further refined using a mask englobing a single tubulin dimer and the density corresponding to single copies of HURP and Kif18A. This density map, which corresponds to an average, clearly shows a steric clash between the expected structural elements from HURP and Kif18A (Figure 4 D). To dissect the different populations that could be contributing to the reconstruction, we proceeded with alignment-free 3D classification, which detected three distinct classes (Figure 4 E). The first class contained density for tubulin and HURP (∼40% of the particles), the second class featured tubulin and Kif18A (∼39% of the particles), while the third class only showed tubulin density (∼21% of the particles) (Supplementary Figure 5 B and Supplementary Figure 8). For the Kif18A-containing class, we were able to build a model (Supplementary Figure 9 A-B) and visualize kinesin-tubulin interactions (Supplementary Figure 9 C-D). Further classification of the Kif18A-containing particles with a mask around the inter-protofilament groove did not show a reliable class where Kif18A displaced HURP’s α-helix but the groove-binding loop was still engaged (not shown). This result further confirms an antagonistic binding mode, where the presence of the motor domain of Kif18A is incompatible with HURP’s MTBD1 engaging the microtubule, since the motor displaced both the α-helix and the groove-binding loop of HURP from tubulin. We concluded that HURP and the motor domain of Kif18A cannot occupy the same tubulin dimer on the microtubule lattice due to steric exclusion.

### HURP and Kif18A synergistically control microtubule length

We next turned our attention to determine how HURP and Kif18A affect microtubule dynamics. Consistent with earlier reports^18–20^, in the absence of free tubulin, 0.1 µM Kif18A led to slow ∼1 nm/s depolymerization of GMPCPP-stabilized microtubules. This depolymerization rate is 4-fold faster than the depolymerization of GMPCPP-microtubules in the absence of Kif18A, but an order of magnitude slower than depolymerization of microtubules by kinesin-13s^38^. The addition of 1 µM HURP^1-173^ mitigated the activity of Kif18A, reducing depolymerization rates to ∼0.05 nm/s (Supplementary Figure 10). We next investigated microtubule dynamics under conditions that included Kif18A with or without HURP. When Kif18A was added to the polymerization mixture alongside free tubulin, the run frequency of motors was significantly reduced (Supplementary Figure 11), likely due to the interaction of Kif18A with free tubulin. To circumvent this issue and ensure attachment of Kif18A to the microtubules, we pre-incubated Kif18A with microtubule seeds in the presence of the slowly hydrolyzable ATP analog AMPPNP. We next flowed a polymerization mixture that included free tubulin, GTP, and ATP into the chamber, allowing us to monitor microtubule growth before and after the dissociation of Kif18A. Upon introduction of ATP in the polymerization mixture, Kif18A started to walk and accumulated at the plus-end of the microtubule (Figure 5 A-B). We noticed a significant decrease in the plus-end growth velocity of microtubules with the accumulation of Kif18A, whereas the minus-end growth remained unaffected. The Kif18A puncta at the plus-end typically released in a single step, and the growth at the plus-end resumed immediately upon Kif18A’s release (Figure 5 A-B). These results are consistent with the idea that Kif18A acts as a ‘molecular cap’ that hinders plus-end growth^26,39^. This capping effect was found to rely on the C-terminal region of Kif18A, since Kif18A^1-480^ failed to exhibit microtubule capping, despite its robust motility on dynamic microtubules (Figure 5 C-D). We propose that the C-terminal MTBD of Kif18A plays a significant role in the plus-end capping effect, either by directly capping the end, or by increasing the dwell time of Kif18A at the microtubule tip.

**Figure 5.**
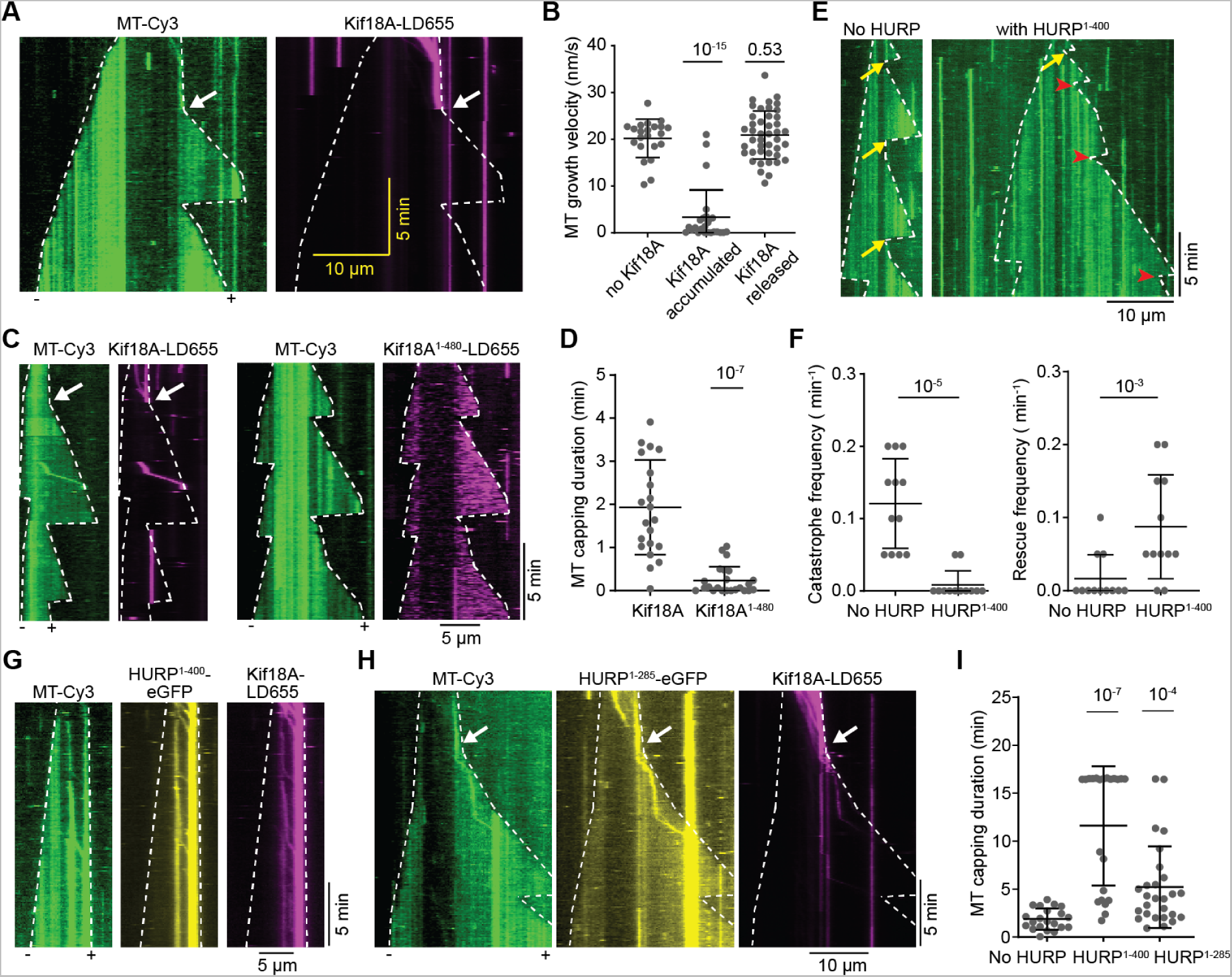
Kif18A and HURP synergistically suppress microtubule dynamics. **A.** Kymographs of dynamic microtubules in the presence of 50 nM Kif18A-LD655 preincubated with GMPCPP-microtubule seeds and AMPPNP (no free Kif18A present during imaging). Upon addition of ATP, Kif18A moves on microtubules and accumulates at the plus-end tip. The plus-end of the microtubule stopped growing until the Kif18A accumulation was spontaneously released (white arrow). **B**. Microtubule plus-end growth velocities with Kif18A accumulated at the plus-end or released from the plus-end (from left to right, n = 22, 25, 43 microtubule growth periods). **C**. (Left) Kymographs of dynamic microtubules with Kif18A-LD655 preincubated on GMPCPP-microtubule seeds and AMPPNP, imaged in the presence of free Kif18A. The white arrow shows the release of Kif18A capping from microtubule plus-end. (Right) Kymographs of dynamic microtubules with Kif18A^1-480^-LD655 preincubated with GMPCPP-microtubule seeds and AMPPNP, imaged in the presence of free Kif18A^1-480^. **D**. Duration of the microtubule plus-end capping by Kif18A or Kif18A^1-480^ (from left to right, n = 21, 23 kymographs). **E.** Kymographs of dynamic microtubules with or without HURP^1-400^. Yellow arrows represent catastrophe events and red arrowheads represent rescue events. **F.** Catastrophe frequencies (left) and rescue frequencies (right) with or without HURP^1-400^ (n = 12 kymographs for each condition) **G.** Kymographs of dynamic microtubules with Kif18A-LD655 and HURP^1-400^-eGFP preincubated with GMPCPP-microtubule seeds and AMPPNP, imaged in the presence of free Kif18A and HURP^1-400^. **H.** Kymographs of dynamic microtubules with Kif18A-LD655 and HURP^1-285^-eGFP preincubated with GMPCPP-microtubule seeds and AMPPNP, imaged in the presence of free Kif18A and HURP^1-285^. The white arrow shows the release of Kif18A capping from the microtubule plus-end. **I**. Duration of the microtubule plus-end capping by Kif18A in conditions of no HURP, 1 μM HURP^1-400^ or 1 μM HURP^1-285^ (from left to right, n = 21, 22 and 27 kymographs). In **A**, **C**, **E**, **G** and **H**, white dashed lines show the track of microtubule ends. In **B**, **D**, **F** and **I**, the center line and whiskers represent the mean and S.D., respectively. P values are calculated from a two-tailed t test. Each imaging duration is 1000s.

Consistent with previous reports, additon of HURP^1-400^ to dynamic microtubules reduced microtubule catastrophes and increased the rescue frequency, without affecting microtubule growth or shrinking rates (Figure 5 E-F, Supplementary Figure 12)^16^. This result led us to investigate HURP’s possible effect on the capping mechanism exerted by Kif18A. The introduction of HURP^1-400^ prolonged Kif18A’s retention at the plus-end, thereby extending the duration at which the Kif18A cap inhibits the growth of the plus-end (Figure 5 G, I). This result is consistent with our findings that HURP^1-400^ prolongs Kif18A’s motility on microtubules (Figure 1 E-F). We also found that HURP^1-400^ co-migrated with Kif18A and accumulated at the plus-end with Kif18A (Figure 5 G). Although HURP^1-285^ also extended Kif18A’s capping period (Figure 5 H-I), the effect was less pronounced, as it did not significantly increase the run time of Kif18A on microtubules (Figure 1 E-F, Figure 2 C-D). Collectively, our findings indicate that HURP and Kif18A synergistically regulate microtubule length. Plus-end accumulation of Kif18A arrests microtubule growth. Kif18A also gradually and slowly depolymerizes stabilized microtubules, but this effect is reduced by the stabilizing activity of HURP. Additionally, Kif18A caps the microtubule plus-end against shrinkage, and this capping activity is enhanced by HURP. Our results show that the combination of HURP and Kif18A substantially suppresses the plus-end dynamics of microtubules and thus maintains a constant microtubule length (Supplementary Figure 13).

## Discussion

In this work, we show that HURP can recruit and activate kinesin-8 Kif18A, providing new insight into how these two proteins organize K-fibers and control their length. Our observations indicate that HURP concentration can be fine-tuned to produce different outcomes on Kif18A motility. At physiologically relevant regimes (i.e. 0.32 µM HURP was reported in *Xenopus laevis* egg extracts^40^), HURP activates Kif18A, presumably by releasing an auto-inhibitory interaction between the motor domain and the C-terminal tail of kinesin. This autoinhibitory mechanism is conserved in other kinesin families, including kinesin-1, kinesin-3, and kinesin-7, and provides a way to locally regulate the engagement of these motors with microtubules ^30–34^.

Our findings suggest that HURP could interact with Kif18A through two distinct sites: The 1-173 segment of HURP, which we refer to as the Kif18A-activating domain, interacts with the motor’s tail, releases auto-inhibition and activates Kif18A motility. In comparison, the 285-400 segment of HURP markedly reduces Kif18A’s velocity, which may be due to an interaction between this region and the 1-480 segment of Kif18A. Interestingly, a cross-talk between the N and C-termini of HURP has been described in previous studies, showing that HURP phosphorylation in the C-terminus by the Aurora A kinase can regulate accessibility of its N-terminus and contribute to HURP localization^29,41,42^. Therefore, the C-terminus of HURP may also regulate the activating role of the HURP N-terminus, resulting in fewer and slower Kif18A runs on the microtubule. Future studies will be required to address the roles of these two regions of HURP, and their post-translational modifications, on the motility of mitotic kinesins.

HURP-mediated activation could promote the enrichment of Kif18A in chromatin-proximal regions of the K-fibers, where HURP localizes during metaphase forming a comet-like gradient^15,42^. Yet, due to saturation of the binding sites by HURP, we observed that increased recruitment of motors to microtubules does not lead to productive motility at higher HURP concentrations. We previously reported a similar regulatory role of MAP7 in kinesin-1 motility^43^, underscoring that concentration-dependent regulation of motors by MAPs could serve as a general mechanism for spatial and temporal regulation of microtubule-driven processes.

The biphasic regulation of Kif18A by HURP concentration we observed *in vitro* can explain why the phenotypes for HURP depletion and overexpression resemble each other *in vivo*. When HURP is depleted, Kif18A likely remains in an auto-inhibited state that limits the number of landing events that result in processive motility. On the other hand, when HURP is overexpressed in pathological states of the cell^12,44–46^, it could saturate the microtubule surface and block efficient Kif18A walking. Both of these situations would lead to inefficient accumulation of Kif18A at the kinetochore-proximal end, promoting defects in chromosome congression during mitosis (Figure 6 A).

**Figure 6.**
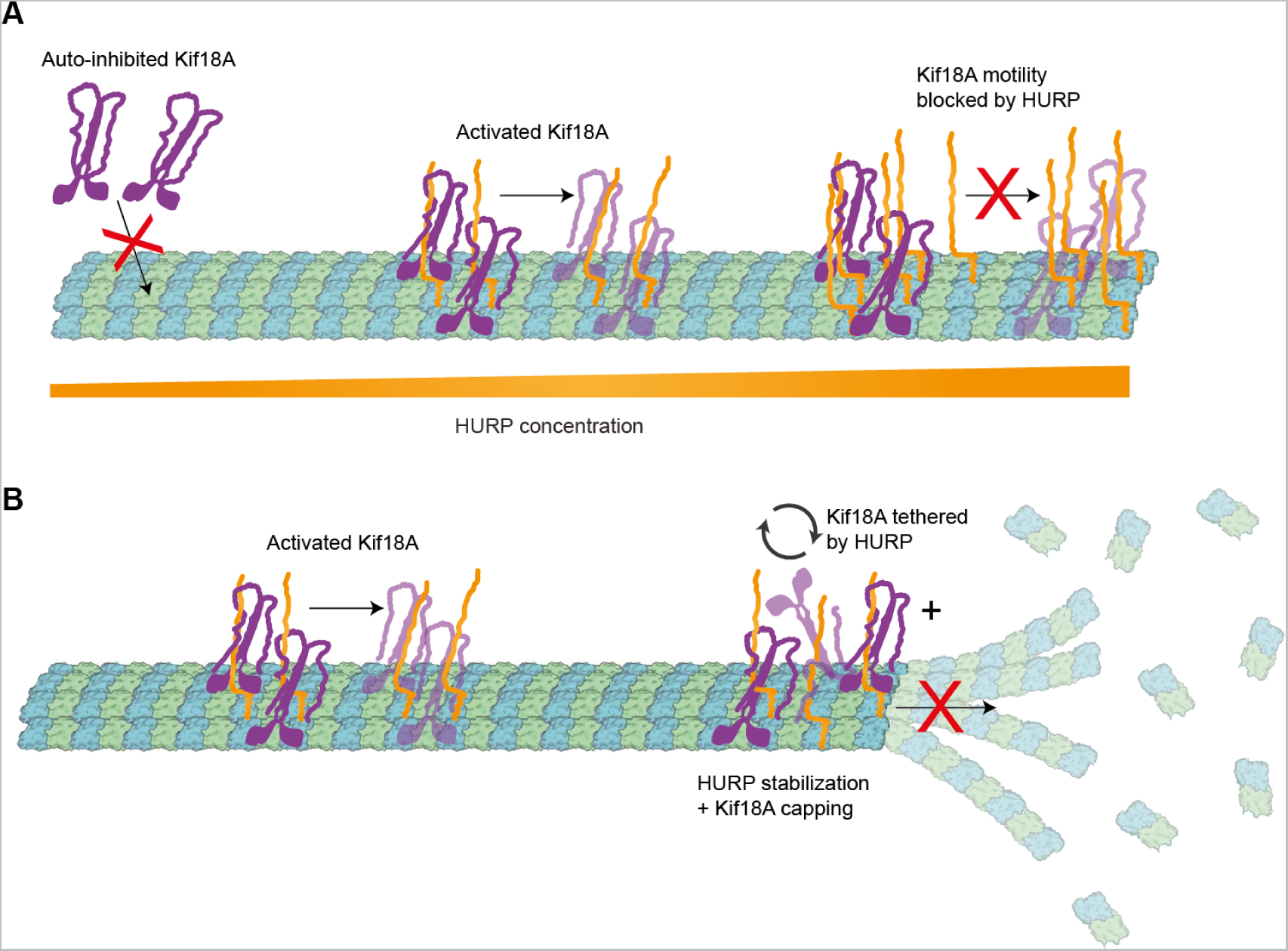
Molecular model of HURP and Kif18A interplay in motility and microtubule dynamics. **A.** Schematic representation of the concentration-dependent regulation of Kif18A motility by HURP. Microtubule-bound HURP helps recruit Kif18A and releases its auto-inhibition. However, excess HURP decreases Kif18A run time and velocity. **B.** Schematic representation of the regulation of microtubule dynamics at the plus-end by HURP and Kif18A. The presence of HURP and Kif18A limits both growth and catastrophe of the microtubule plus-end, regulating microtubule length.

In kinesin-8 Kip3p, the yeast homolog of Kif18A, the length-dependent accumulation on the plus-end of K-fibers has been attributed to kinesins randomly landing on the microtubule surface and processively walking towards the kinetochore-proximal end, where they would accumulate and quench polymerization dynamics. This “antenna model” explained how members of the kinesin-8 family could depolymerize longer microtubules at a faster rate than shorter ones^22,25,47^. We suggest that productive landing of Kif18A might not occur randomly *in vivo*, but instead be more prevalent in regions where HURP localizes and activates the motor, closer to the plus-end of K-microtubules, thus facilitating its accumulation.

Cryo-EM imaging of microtubule-bound HURP revealed how this MAP associates with microtubules through a bipartite binding motif that stabilizes adjacent tubulin dimers by establishing lateral contacts. Consistent with its role as a stabilizer, HURP has been implicated in attenuating microtubule dynamic instability^48^ by reducing the catastrophe frequency and increasing the rescue frequency *in vitro*^16^. Our structural model suggests that this function could be exerted by laterally stabilizing protofilaments, and preventing their peeling at the microtubule tip during catastrophes^49,50^.

The microtubule binding mechanism of HURP resembles that of another SAF and stabilizer, TPX2^36,51,52^. Both HURP and TPX2 are regulated by the Ran-GTP pathway^14,53^, both interact with mitotic kinesins (HURP with Kif18A and Kif11/Eg5^17,42^ and TPX2 with Kif11/Eg5 and Kif15^54–56^), and both nucleate and stabilize K-microtubules^14,41,57–60^. The structured binding motif in HURP stabilizes lateral interactions, while TPX2 establishes both lateral and longitudinal contacts. Intriguingly, TPX2 has been shown to interact with HURP and other partners for bipolar spindle formation^13^, which could point to distinct but potentially complementary roles in assembling K-fibers.

Finally, we demonstrate that Kif18A localizes to the plus-ends of microtubules and caps their growth, likely through a mechanism that relies on the C-terminal tail of the motor. Given the previous identification of a non-motor MTBD at the C-terminal of Kif18A^39,61^, we propose that this region significantly influences the capping process. It remains to be demonstrated whether the C-terminal MTBD directly tethers to the plus-end and blocks the addition of new tubulin units, or, if it facilitates Kif18A’s accumulation and retention at the plus-end by increasing its dwell time.

HURP enhances Kif18A’s capping effect by increasing the residence time of the motor at the microtubule plus-end, in addition to having its own role as a stabilizer to reduce shrinkage. We propose that HURP and Kif18A work synergistically to reduce microtubule dynamics, which could serve as a spindle length control mechanism during mitosis (Figure 6 B). Our observation that HURP can be transported towards the plus-end by Kif18A has been reported for other MAPs^62^ and suggests that this could play a secondary role in creating a HURP gradient, besides the canonical Ran-mediated pathway that dictates HURP distribution. Recent studies have reported that HURP shows differential binding to microtubules of variable length, through a mechanism that is still unclear and could involve centrosomal regulation^63^. Future studies will be required to reveal how HURP “senses” K-fiber length and tune its dynamics together with Kif18A.

## Materials and Methods

### Protein expression, purification and labeling

HURP constructs were cloned from U2OS-derived cDNA and either inserted into a pRSFDuet-SUMO vector for bacterial expression, or into a TwinStrep-pFastBac vector for insect cell expression through a sequence and ligase independent cloning (SLIC) strategy. Kif18A constructs were produced from the pMX229 Addgene plasmid deposited by Linda Wordeman. Linear inserts containing Kif18A residues 1-373, 1-480, and 1-898 were PCR-cloned from pMX229 and inserted into a pRSFDuet-SUMO vector carrying the coding sequence for a C-terminal SNAP tag as described above. The sequence of all constructs was verified either by Sanger or full-length plasmid sequencing.

Kif18A (full-length Kif18A-SNAP, Kif18A^1-480^-SNAP) and truncated HURP constructs (HURP^1-173^-eGFP, HURP^1-285^, HURP^1-285^-eGFP, HURP^1-400^-eGFP) were transformed to Rosetta2(DE3) competent cells, plated for kanamycin selection, and a single colony was grown in LB+kanamycin at 37°C until OD reached 0.6. Expression was induced with 0.2 mM IPTG at 37 °C and cells were harvested after 4 h. Full-length HURP^1-846^-eGFP was produced in Sf9 cells through baculovirus infection as previously described^16^.

Cell pellets of HURP^1-173^-eGFP, HURP^1-285^, HURP^1-285^-eGFP, HURP^1-400^-eGFP were lysed through sonication in His lysis buffer (20 mM HEPES pH 7.5, 300 mM KCl, 1 mM MgCl_2_, 5% glycerol, 20 mM imidazole, 0.1% Tween-20, 10 mM BME, 1x benzonase, 1 protease inhibitor tablet) and the lysate was cleared by centrifugation at 18,000 rcf for 1 h at 4 °C. The cleared lysate was incubated with 2 mL of Ni-NTA agarose resin, previously equilibrated in lysis buffer, for 2 h at 4°C. The resin was washed with 150 mL buffer, alternating washing buffer (20 mM HEPES pH 7.5, 300 mM KCl, 1 mM MgCl_2_, 5% glycerol, 20 mM imidazole, 0.1% Tween-20, 10 mM BME) with washing buffer supplemented with 700 mM NaCl. The protein was then eluted overnight in washing buffer supplemented with the Ulp1 protease to cleave the N-terminal His-SUMO tag on the constructs. The eluted protein was diluted 4x with IEX A buffer (20 mM HEPES pH 7.5, 100 mM NaCl, 10 mM BME) and loaded onto a 5 mL HiTrap SP HP column for ion exchange chromatography. The protein was then eluted with a salt gradient from 0 to 50% IEX B buffer (20 mM HEPES pH 7.5, 2 M NaCl, 10 mM BME) and HURP-containing fractions were pooled, concentrated, and buffer-exchanged to SEC buffer (20 mM HEPES pH 7.5, 300 mM KCl, 1 mM MgCl_2_, 5% glycerol, 1 mM TCEP). The concentrated sample was loaded onto a Superdex 200 10/300 GL column, and eluted with 1.2 CV of SEC buffer. Samples containing the protein of interest were pooled, concentrated, aliquoted and flash frozen in liquid nitrogen for storage at −80°C.

HURP^1-846^-eGFP was purified as previously described^16^. Briefly, after cell lysis and centrifugation, the cleared lysate was filtered through a 0.2 μM filter and loaded onto a 5 mL HiTrap SP HP column. After washing, the protein was eluted with a salt gradient ranging from 240 mM to 1 M NaCl. Protein-containing fractions were concentrated, diluted to lower salt and loaded onto a 1 mL HiTrap Q HP column. The flow-through was collected, pooled, and loaded onto a 5 mL HisTrap HP. HURP was eluted with an imidazole gradient and buffer exchanged to the final storage buffer E (50 mM HEPES pH 8.0, 300 mM KCl, 1 mM DTT). The concentrated sample was injected onto a Superdex 200 10/300 GL column. Fractions containing HURP^1-846^-eGFP were pooled, concentrated, aliquoted, and flash-frozen in liquid nitrogen for storage at −80°C.

Kif18A SNAP constructs were purified as follows. Cell pellets were resuspended, lysed in Kif18A buffer (25 mM Tris pH 7.5, 300 mM KCl, 5 mM MgCl_2_, 20 mM imidazole, 0.1% Tween-20, 1 mM ATP, 1 mM EGTA, 1 mM DTT, 5% glycerol), and centrifuged as described for the other proteins. The cleared lysates were incubated with 2 mL Ni-NTA agarose resin and equilibrated in the Kif18A buffer for 2 h. The resin was washed four times with 30 mL Kif18A buffer. After washing, the agarose resin was resuspended in 8 mL Kif18A buffer supplemented with Ulp1 protease. Elution with Ulp1 was carried overnight at 4°C, after which the protein was completely released from the Ni-NTA beads. Beads were separated from the solution by gentle centrifugation and the solution was concentrated to 800 μL. The SNAP tag was labeled with LD655 by incubating 15 nmoles of LD-655 benzylguanine (Lumidyne) with concentrated protein solution for 5 h at 4°C. The mixture was then centrifuged to remove any aggregates, and injected onto a Superdex 200 10/300 GL gel filtration column. Protein-containing fractions were pooled, concentrated, aliquoted and frozen before storing them at −80°C.

### Cryo-EM sample preparation

Porcine brain tubulin (Cytoskeleton Cat # T240) was reconstituted to 10 mg/mL in BRB80 buffer (80 mM Pipes, pH 6.9, 1 mM ethylene glycol tetraacetic acid (EGTA), 1 mM MgCl_2_) with 10% (v/v) glycerol, 1 mM GTP, and 1 mM DTT. 10 μL of the tubulin solution were polymerized at 37°C for 15 min. 1 μL of 2 mM taxol was added to the polymerizing tubulin, and incubated at 37°C for 10 minutes; this was followed by a second addition of 1 μL taxol and a further incubation of 30 minutes. microtubules were pelleted by centrifugation at 37°C and 15,000 rcf for 20 minutes. The supernatant containing free tubulin was discarded and the pelleted microtubules were resuspended in resuspension buffer (BRB80 buffer supplemented with 0.05% NP-40, 1.5 mM MgCl_2_, 1 mM DTT and 250 μM taxol). After measuring the tubulin concentration in a CaCl_2_ depolymerised aliquot, the microtubule solution was diluted to 2 μM in dilution buffer (BRB80 buffer supplemented with 0.05% NP-40, 1.5 mM MgCl_2_, 1 mM DTT and 100 μM taxol). Immediately before sample preparation, all microtubule-binding proteins were desalted to cryo buffer (BRB80 buffer supplemented with 0.05% NP-40, 1.5 mM MgCl_2_, 1 mM DTT) using Zeba Spin desalting columns (Pierce).

To prepare microtubule-HURP^1-285^ samples, 2 μL of 2 μM taxol-microtubules were incubated on a glow-discharged holey carbon cryo-EM grid (QuantiFoil, Cu 300 R 2/1) for 30 seconds, manually blotted with Whatman filter paper, and 2.5 μL of 30 μM HURP^1-285^ were added to the grid. The grid was transferred to a Vitrobot (Thermo Fisher Scientific) set at 25 °C and 80% humidity, and plunge-frozen in liquid ethane after a 1 minute incubation with a blot force of 6 pN and a blot time of 6 s.

Microtubule-HURP^1-285^-eGFP-Kif18A^1-373^-SNAP-LD655 sample preparation followed a similar procedure. 2 μL of 2 μM taxol-microtubules were incubated on a glow-discharged holey carbon cryo-EM grid (QuantiFoil, Au 300 R 1.2/1.3) for 30 seconds, manually blotted as before, and incubated with 2.5 μL of a mixture with 8 μM HURP^1-285^-eGFP, 8 μM Kif18A^1-373^-SNAP-LD655, and 5 mM AMPPNP. After incubating for 1 minute in the Vitrobot under identical conditions as the previous sample, the grid was plunge-frozen and transferred to liquid nitrogen as described before.

### Cryo-EM data collection

Data for microtubules decorated with HURP^1-285^ and HURP^1-285^ eGFP + Kif18A^1-373^-SNAP-LD655 were collected using an Arctica microscope (Thermo Fisher Scientific), operated at an accelerating voltage of 200 kV (Table S1). All cryo-EM images were acquired on a K3 direct electron detector (Gatan), at a nominal magnification of 36,000×, corresponding to a calibrated physical pixel size of 1.14 Å. The camera was operated in superresolution mode, with a dose rate of ∼7.2 electrons/pixel/s on the detector. We used an exposure time of ∼9 s dose-fractionated into 50 frames, corresponding to a total dose of ∼50 electrons/Å^2^ on the specimen. All the data was collected semi-automatically with the SerialEM software package^64^.

### Cryo-EM image processing

For the microtubule-HURP^1-285^ dataset, the movie stacks were imported to CryoSparc and motion-corrected^65^. The CTF parameters were estimated with the patch CTF job and manually curated to remove bad micrographs. Particles were automatically picked with the filament tracer, initially without a reference, and later using 2D templates as input. The segment separation was set to 82 Å, corresponding to the length of an α-𝛽 tubulin dimer. Particle images were extracted with a box size of 512 pixels and Fourier-cropped to 256 pixels for initial image processing. These images were subjected to 2 rounds of 2D classification and classes showing clear density for the microtubule were selected; classes showing blurry density, junk particles or non-centered microtubules were discarded. Microtubules with different numbers of protofilaments were separated through a heterogeneous refinement job where 13 and 14 PF lowpass-filtered references were used as initial models. In both cryo-EM datasets, the 14 PF population corresponded to the majority class, and therefore was selected for further processing. 14 PF particles were subjected to a helical refinement with an initial rise estimate of 82.5 Å and a twist of 0°. Angular assignments and shifts were further refined through a local refinement using a hollow cylindrical mask enclosing the microtubule reconstruction. To obtain a reconstruction that accounts for the presence of a symmetry-breaking seam on the microtubule, we used a Frealign-based seam search routine with custom scripts that determines the seam position on a per-particle basis^35^. For this purpose, we converted the CryoSparc alignment file from the last local refinement, first to the STAR format using the csparc2star script from the PyEM suite (10.5281/zenodo.3576630), and then to PAR Frealign format using a custom

Python script. Upon completion of the seam search protocol, the particles were imported back to CryoSparc with the seam-corrected alignments and re-extracted without Fourier cropping (box size: 512 pixels), using the new improved alignments to recenter the picks before extraction. A volume was reconstructed with the imported particles through a local refinement job and a local CTF refinement was performed to estimate the per-particle CTF. Another local refinement was run on this particle set to produce the final C1 reconstruction. The symmetry search job was used to determine the rise and the twist of the map, and these parameters were input to a symmetry expansion job to fully exploit the pseudo-symmetry of each microtubule particle. A local refinement was performed on the new expanded particle stack with a cylindrical mask around the microtubule to yield the final symmetrized reconstruction of the entire microtubule. A final local refinement was performed with a smaller mask around the good PF (PF opposite to the seam) and the PF adjacent to it, producing a 3.1 Å resolution map of HURP bound to microtubules (Supplementary Figure 4, Supplementary Figure 5 A).

The HURP^1-285^ eGFP + Kif18A^1-373^-SNAP-LD655 dataset was processed in an identical way up to the step where the particle set is symmetry expanded and the final symmetrized reconstruction of the entire microtubule is generated. From this point, we generated a new local refinement with a shaped mask encompassing 2 tubulin dimers, 2 Kif18A monomers, and the inter-PF groove where HURP inserts. This produced a consensus reconstruction at 2.9 Å resolution that was further sorted using 3D classification, with a more constrained map only covering single copies of a tubulin dimer, a Kif18A, and HURP molecule. Initially, a random subset containing 10% of the refined particles was used for reference-free 3D classification without alignment, using the consensus reconstruction and aligned particles as inputs. This generated 5 classes that were used as initial references for a second alignment-free 3D classification job on the full particle stack. Some of these classes contained density for tubulin + HURP, others showed tubulin + Kif18A, and 1 class only showed density for tubulin. The second 3D classification job recapitulated the results from the first classification and allowed us to computationally sort the compositional heterogeneity present in the consensus reconstruction. Classes showing no distinct features with each other were combined and locally refined, yielding a 3.5 Å resolution class for tubulin, a 3 Å resolution map for tubulin + Kif18A, and a 3 Å resolution reconstruction for tubulin + HURP (Supplementary Figure 5 B, Supplementary Figure 8). Cryo-EM processing parameters are shown in Table S1.

### Model building and refinement

The final cryo-EM map from the microtubule-HURP^1-285^ dataset was used for modeling of HURP MTBD1. Tubulin dimers from a previous publication (PDB: 6DPV)^66^ were rigid-body fitted into the density map using ChimeraX^67^. The local resolution of the map region corresponding to HURP and the presence of 3 sequential bulky side-chains in the inter-PF groove HURP density (R122, Y123, R124) allowed us to unambiguously assign the protein sequence register. HURP modeling was performed in Coot^68^ by manually tracing the main chain and assigning the corresponding residues. In the case of the α-helical density, it was modeled as a perfect α-helix and the register was identified by the presence of bulky side chains L94 and Y97. The HURP and tubulin models (2 tubulin dimers, 1 HURP) were then combined in a single PDB file and real-spaced refined using the Phenix software^69^. For microtubule-Kif18A modeling a similar procedure was followed starting from PDB ID 5OCU^37^, manually applying changes and then refining it against the Kif18A-containing density map. Model refinement statistics are shown in Table S1. For visualization purposes and figure generation, refined models for α-β-tubulin, HURP or Kif18A were superimposed with the corresponding maps, colored based on subunit type (α-tubulin in green, β-tubulin in blue, HURP in orange and Kif18A in purple), and the maps were colored and segmented accordingly, creating independent maps for the four subunit types. The splitted maps for the different types of subunits were set to different thresholds that better reflected their average local resolution. Lower and identical threshold values were used for α and β tubulins to highlight their higher resolution features, while the thresholds for HURP and Kif18A were set independently.

### Native tubulin extraction

We extracted native tubulin from pig brain through a series of polymerization and depolymerization cycles. Initially, the pig brain tissue was lysed and then mixed in a 1:1 mass-to-volume ratio with a depolymerization buffer (50 mM MES, 1 mM CaCl_2_, pH 6.6 with NaOH). This mixture was centrifuged at 8,000 rpm and 4 °C. The supernatant was then combined with high molarity PIPES buffer (HMPB) (1M PIPES Free Acid, 10 mM MgCl_2_, and 20 mM EGTA, pH 6.9 with KOH) and glycerol in a 1:1:1 volume ratio. Subsequently, GTP and ATP were added to reach final concentrations of 0.5 mM and 1.5 mM, respectively. This mixture was incubated at 37 °C for 1 hour, followed by centrifugation at 400,000 rcf for 30 minutes at 37 °C.

After this process, the pellet was resuspended in the depolymerization buffer and incubated at 4 °C for 15 minutes to induce depolymerization. The supernatant obtained was then mixed again with HMPB and glycerol in a 1:1:1 volume ratio, with the addition of GTP and ATP to the previously stated concentrations. The mixture underwent another incubation at 37 °C for 1 hour, followed by centrifugation at 400,000 rcf for 30 minutes at 37 °C. The final pellet was resuspended in cold BRB80 buffer and incubated at 4 °C for 30 minutes. A last centrifugation at 400,000 rcf for an unspecified duration at 4 °C was performed, and tubulin was diluted to 34 mg/mL before the tubulin was stored at −80 °C.

### Biotin or Cy3 labeling of tubulin

To label tubulin with biotin or Cy3, NHS ester labeling was performed on polymerized microtubules. Initially, GTP and DTT were added to 0.2 mL of tubulin aliquots, adjusting the concentrations to 5 mM each. This mixture was incubated at 37°C for 30 minutes to promote microtubule polymerization. Following polymerization, the mixture was centrifuged at 400,000 rcf at 37°C for 30 minutes, overlaying it with 0.5 mL of warm high pH cushion (0.1M HEPES, pH 8.6, 1 mM MgCl_2_, 1 mM EGTA, 60% (v/v) glycerol) to enhance pellet separation. The supernatant above the cushion was then discarded, and the interface between the tube wall and cushion was gently washed twice with 250 µL of warm labeling buffer (0.1M HEPES, pH 8.6, 1 mM MgCl_2_, 1 mM EGTA, 40% (v/v) glycerol) before complete removal of the supernatant and cushion. Subsequently, the pellet was resuspended in 0.4 mL of warm labeling buffer, to which 50 µL of 6 mM NHS-biotin or NHS-Cy3 in DMSO was added. This solution was then incubated at 37°C for 30 minutes on a roller mixer.

To eliminate free NHS-biotin or NHS-Cy3, the microtubule mixture underwent centrifugation at 400,000 rcf at 37°C for 30 minutes using a low pH cushion (BRB80 with 60% (v/v) glycerol). The supernatant above the cushion was removed, and the tube wall-cushion interface was rinsed twice with 250 µL of warm BRB80 before discarding the supernatant and cushion. The tube wall was then washed again twice with 250 µL of warm BRB80. The pellet was resuspended in cold BRB80 and incubated at 4°C for 30 minutes to allow microtubule depolymerization. Finally, the mixture was centrifuged at 400,000 rcf at 4°C for 30 minutes, and the supernatant was collected and stored in a −80°C freezer.

### Preparation of GMPCPP-microtubule seeds

GMPCPP-microtubule seeds were used for the dynamic microtubule assays. The ultracentrifuge, rotor, tubes, and BRB80 buffer (80 mM PIPES (Free Acids), 1 mM MgCl_2_, 1 mM EGTA, 1 mM DTT, pH adjusted to 6.8 with KOH, ensuring a pH below 7) with 10% DMSO were pre-cooled on ice. A mixture was then made with unlabeled tubulin, 5% biotin-tubulin, and 5% Cy3-labeled tubulin (Cy3-labeled tubulin is optional). The mixture was diluted to 1 to 3 mg/mL with cold BRB80 containing 10% DMSO and incubated on ice for 10 minutes (for longer microtubules, a concentration of 0.3 to 0.5 mg/mL was used). The mixture was cold spun at 400,000 rcf for 10 minutes to remove inactive tubulin, and the supernatant was collected, to which GMPCPP was added to achieve a final concentration of 1 mM before incubation at 37°C for 20 minutes (extended to 90 to 120 minutes for longer microtubules). After warming the centrifuge equipment and buffer to 37°C, the tubulin mix was spun at 37°C at 400,000 rcf for 10 minutes. The supernatant was discarded, and the pellet was gently resuspended in 25 to 50 µL of warm BRB80 buffer (using a cut pipette tip to avoid breaking the microtubules). The GMPCPP microtubule seeds were usually good for 2 weeks.

### Preparation of taxol-stabilized microtubules for TIRF

Taxol stabilized microtubules were used for single molecule motility imaging. They were made by diluting 4 μL of 34 mg/mL of tubulin, 5% of which was biotin-labeled and 5% which was Cy3-labeled, into 46 μL BRB80 ( 80 mM PIPES at pH 6.8, 1 mM MgCl_2_, and 1 mM EGTA). This mixture was then added to an equal volume of polymerization solution (1X BRB80 with 2 mM GTP and 20% DMSO). The tubulin was incubated at 37°C for polymerization for 40 minutes, after which 10 nM of taxol was added and the mixture was incubated for another 40 minutes. The microtubules were pelleted by centrifugation at 20,000 rcf for 15 minutes at 37°C and then resuspended in 25 μL BRB80 solution with 10 nM taxol and 1 mM DTT. The microtubule is stocked at room temperature and good to use for 2 weeks.

### Fluorescence microscopy

Fluorescence imaging utilized a custom-built multicolor objective-type TIRF microscope, incorporating a Nikon Ti-E microscope body, a 100X magnification 1.49 N.A. apochromatic oil-immersion objective (Nikon), and a Perfect Focus System. Detection of fluorescence employed an electron-multiplied charge-coupled device camera (Andor, Ixon EM+, 512 × 512 pixels), with an effective camera pixel size of 160 nm post-magnification. Excitation of GFP, Cy3, and LD655 probes occurred via 488 nm, 561 nm, and 633 nm laser beams (Coherent) delivered through a single mode fiber (Oz Optics), with emission filtering accomplished using 525/40, 585/40, and 697/75 bandpass filters (Semrock), respectively. Microscope operations were managed via MicroManager 1.4.

### Preparation of flow chambers

Glass coverslips were coated s with polyethylene glycol (PEG) to reduce nonspecific protein binding. First, plain glass coverslips underwent sequential cleaning steps involving water, acetone, and water sonication for 10 min each, followed by a 40 min sonication in 1 M KOH using a bath sonicator. Subsequently, the coverslips were rinsed with water, immersed in 3-Aminopropyltriethoxysilane in acetate and methanol for 10 min with 1-min sonication intervals between steps, further cleaned with methanol, and air-dried. A 30 µl volume of 25% biotin-PEG-succinimidyl valerate in a NaHCO_3_ buffer (pH 7.4) was applied between two coverslip pieces and left to incubate at 4 °C overnight. Following incubation, the coverslips were cleaned with water, air-dried, vacuum sealed, and long-term stored at −20 °C. Flow chambers were constructed by sandwiching double-sided tape between a PEG-coated coverslip and a glass slide. To facilitate solution flow into the chamber during real-time DDR motility recording, two holes were drilled at each end of the chamber on the glass slides.

### Single-molecule motility imaging

The flow chambers underwent a 2-minute incubation with 5 mg/ml streptavidin, followed by washing with MB buffer (composed of 30 mM HEPES pH 7.0, 5 mM MgSO_4_, 1 mM EGTA, 1 mg/ml casein, 0.5% pluronic acid, 0.5 mM DTT, and 1 µM Taxol). Subsequently, the chamber was incubated with biotinylated microtubules for 2 minutes and washed again with MB buffer. Proteins were then diluted to desired concentrations in imaging buffer (MB buffer supplemented with 0.15 mg/ml glucose oxidase, 0.025 mg/ml catalase, 0.8% D-glucose, and 2 mM ATP), and introduced into the flow chamber. Motility was recorded over a 2-minute period.

### Dynamic microtubule imaging

To conduct a dynamic microtubule assay, tubulin, HURP, and Kif18A were cold centrifuged at 400,000 rcf for 10 minutes to eliminate protein aggregates. Subsequently, biotinylated GMPCPP-microtubule seeds (with or without Cy3-labeling) were incubated on a biotin-PEG glass surface via Streptavidin bonding. The chamber was then flowed with a dynamic microtubule mixture, (1X BRB80 pH 6.8, 1 mg/ml casein, 0.5% pluronic acid, 0.5 mM DTT, 150 mM KAc, 2 mg/mL unlabeled tubulin, 0.05 mg/mL Cy3-tubulin, 4 mM GTP, 1 mM ATP, and 0.2% methyl-cellulose), along with the desired concentrations of Kif18A and/or HURP.

For the pre-incubation of Kif18A on the microtubule seeds, Kif18A was pre-incubated in MB buffer containing 150 mM KAc and AMPPNP for 30 minutes to enhance its affinity to microtubule.

## Acknowledgements

We thank J. Peukes and Z. Yang for helpful discussions related to image processing and biochemical design, L. Wordeman for Kif18A plasmid Addgene deposition, M. Esbin for supplying the U2OS cells used in cloning experiments, D. Toso, R. Thakkar, P. Tobias, and K. Stine for their support with cryo-EM data collection and computation infrastructure, the UCSF ChimeraX team for the software development used in structural rendering, V. Perez-Bertoldi for assisting with graphic design and illustration, Y. He for producing the pegylated glass surface used in TIRF microscopy and J. Fernandes for tubulin preparation. Work was funded by NIGMS (R35GM127018 to E.N., R35GM136414 to A.Y.), by the European Research Council (ERC-2022-SYG to E.N.) and by NSF (MCB-1055017, MCB-1617028 to A.Y.). E.N. is a Howard Hughes Medical Institute Investigator.

## Author contributions

J.M.P.B., A.T. and E.N. initially conceived the project and all authors contributed to its development and the design of experiments. J.M.P.B purified proteins, performed cryo-EM experiments, analyzed data and performed modeling. Y.Z. performed single-molecule TIRF experiments, dynamic microtubule imaging and analyzed data.

E.N and A.Y. supervised the project and secured funding. J.M.P.B., Y.Z., E.N. and A.Y. wrote the manuscript, with further edits by all authors.

## Supplementary Figures and Tables

**Supplementary Figure 1.**
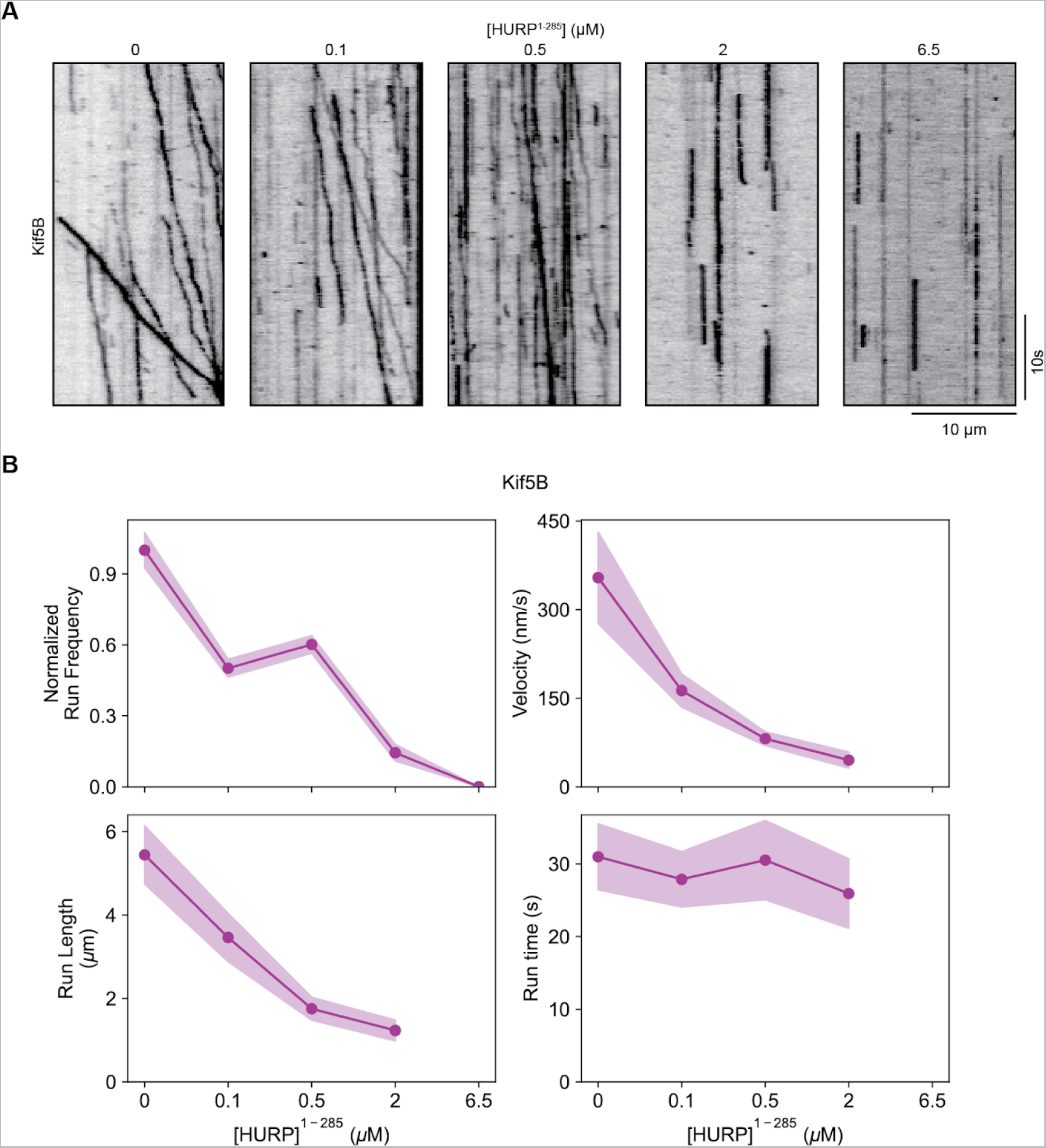
HURP fails to activate Kif5B motility. **A**. Representative kymographs showing the motility of full-length Kif5B in the presence of increasing HURP^1-285^. **B.** Normalized run frequency (n = 10 kymographs for each condition), velocity, run length and run time (from left to right, n = 25 motors for each condition) of Kif5B for different HURP^1-285^ concentrations. The line and shadows represent the mean and S.E., respectively.

**Supplementary Figure 2.**
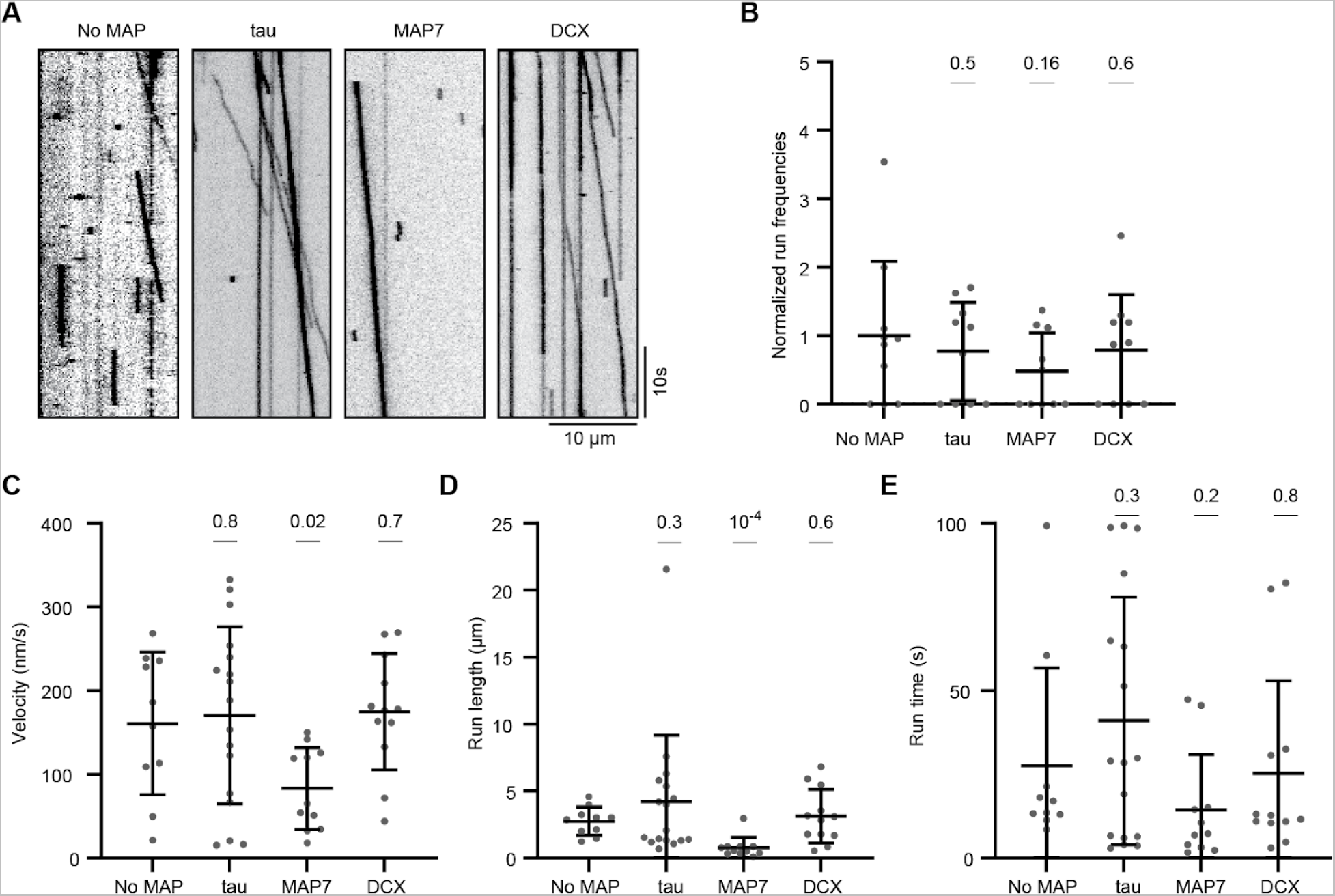
Tau, MAP7 and DCX fail to activate Kif18A motility. **A**. Representative kymographs showing motility of Kif18A in the presence of tau, MAP7 and DCX. **B -E.** Normalized run frequency (**B**), velocity (**C**), run length (**D**) and run time (**E**) of Kif18A in the presence of tau, MAP7 or DCX (For **B**, n = 10 kymographs for each condition; for **C** - **E**, n = 10, 17, 11, 12 motors). The center line and whiskers represent the mean and S.D., respectively. P values are calculated from a two-tailed t test.

**Supplementary Figure 3.**
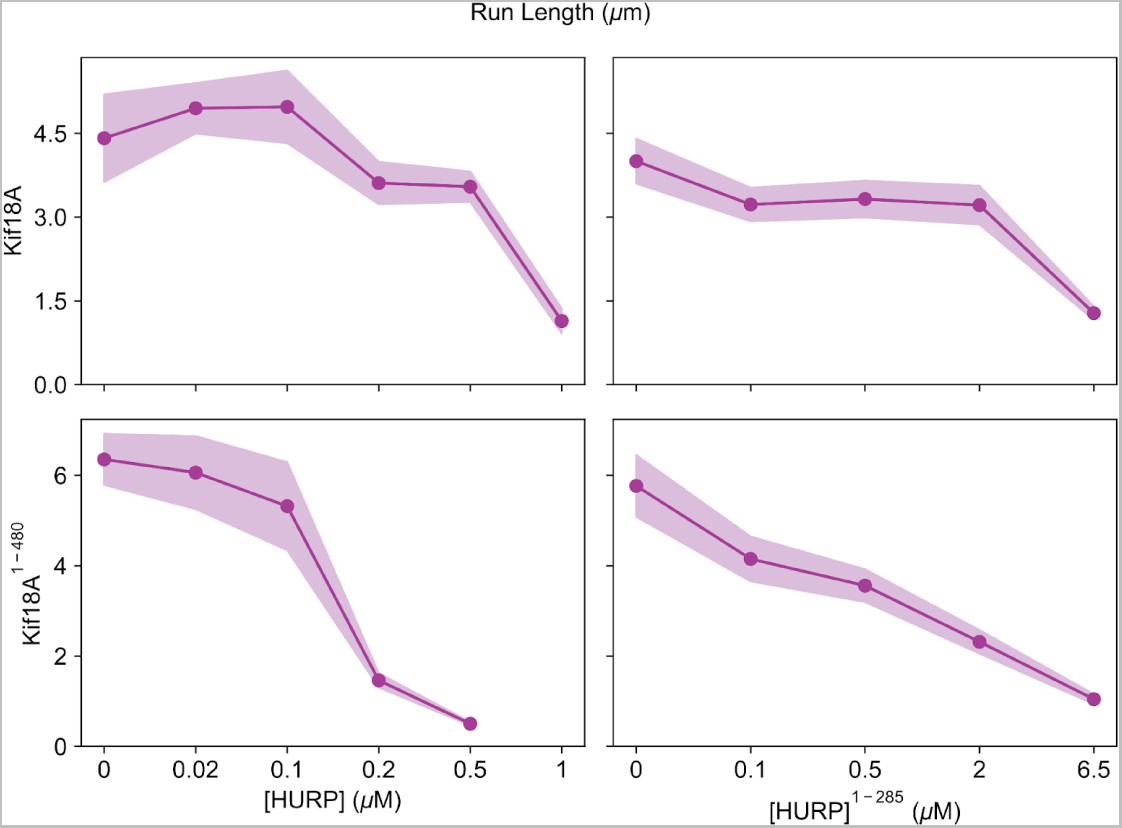
High concentrations of HURP restrict the run length of Kif18A. **A**. Run length of Kif18A with titrated HURP (upper left, n = 32, 50, 33, 48, 50, 40 motors, respectively), Kif18A with titrated HURP^1-285^ (upper right, n = 51, 44, 54, 52, 52 motors, respectively), Kif18A^1-480^ with titrated HURP (lower left, n = 25 motor for each data point) and Kif18A^1-480^ with titrated HURP^1-285^ (lower right, n = 52, 51, 52, 52, 52 motors, respectively). The line and shadows represent the mean and S.E., respectively.

**Supplementary Figure 4.**
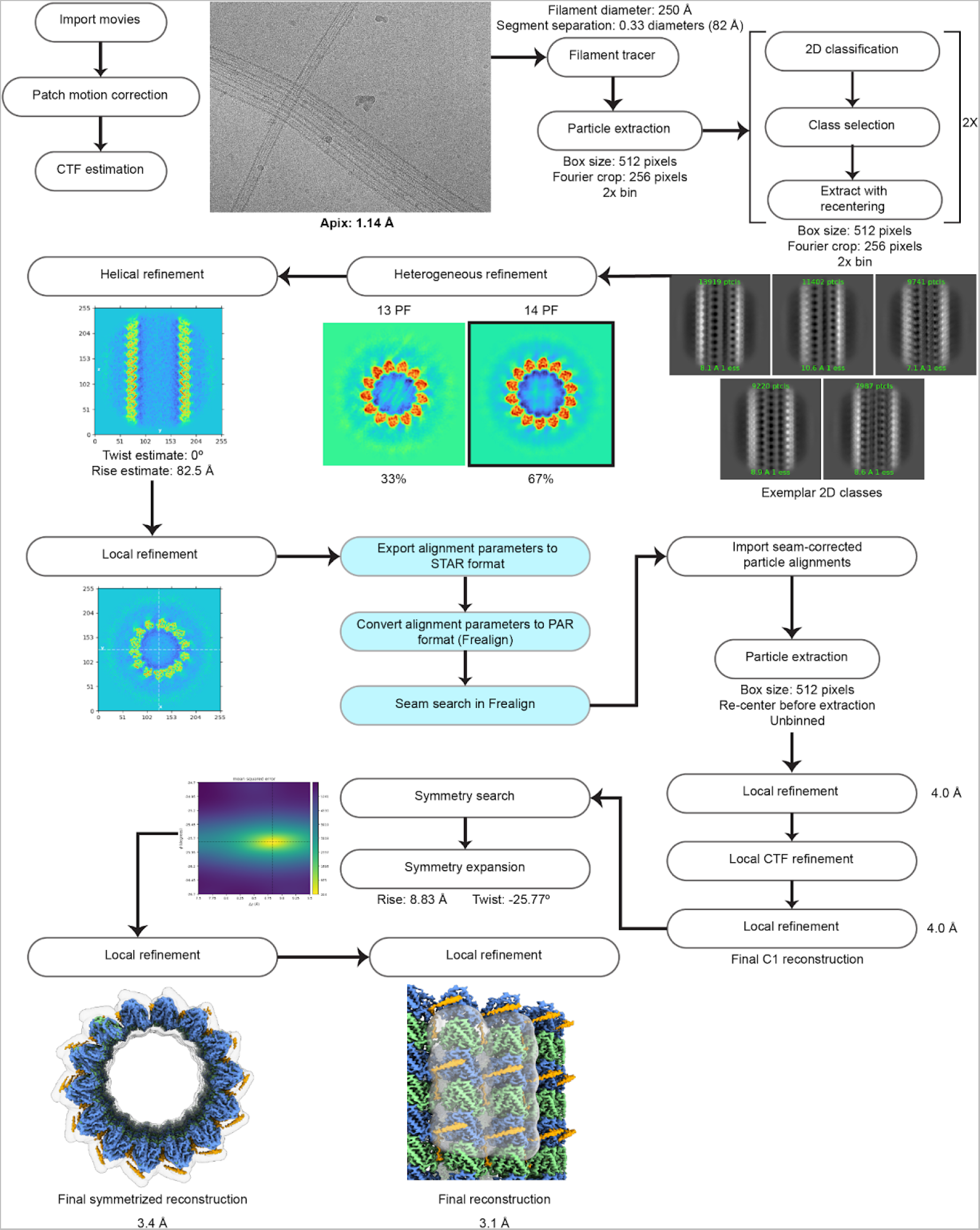
Cryo-EM data processing for the microtubule-HURP dataset. Processing pipeline applied to the HURP-bound microtubule dataset. Unless specified otherwise, all steps were performed in CryoSparc. Boxes in light blue include steps implemented outside of the CryoSparc software package. Masks are shown with transparent surfaces.

**Supplementary Figure 5.**
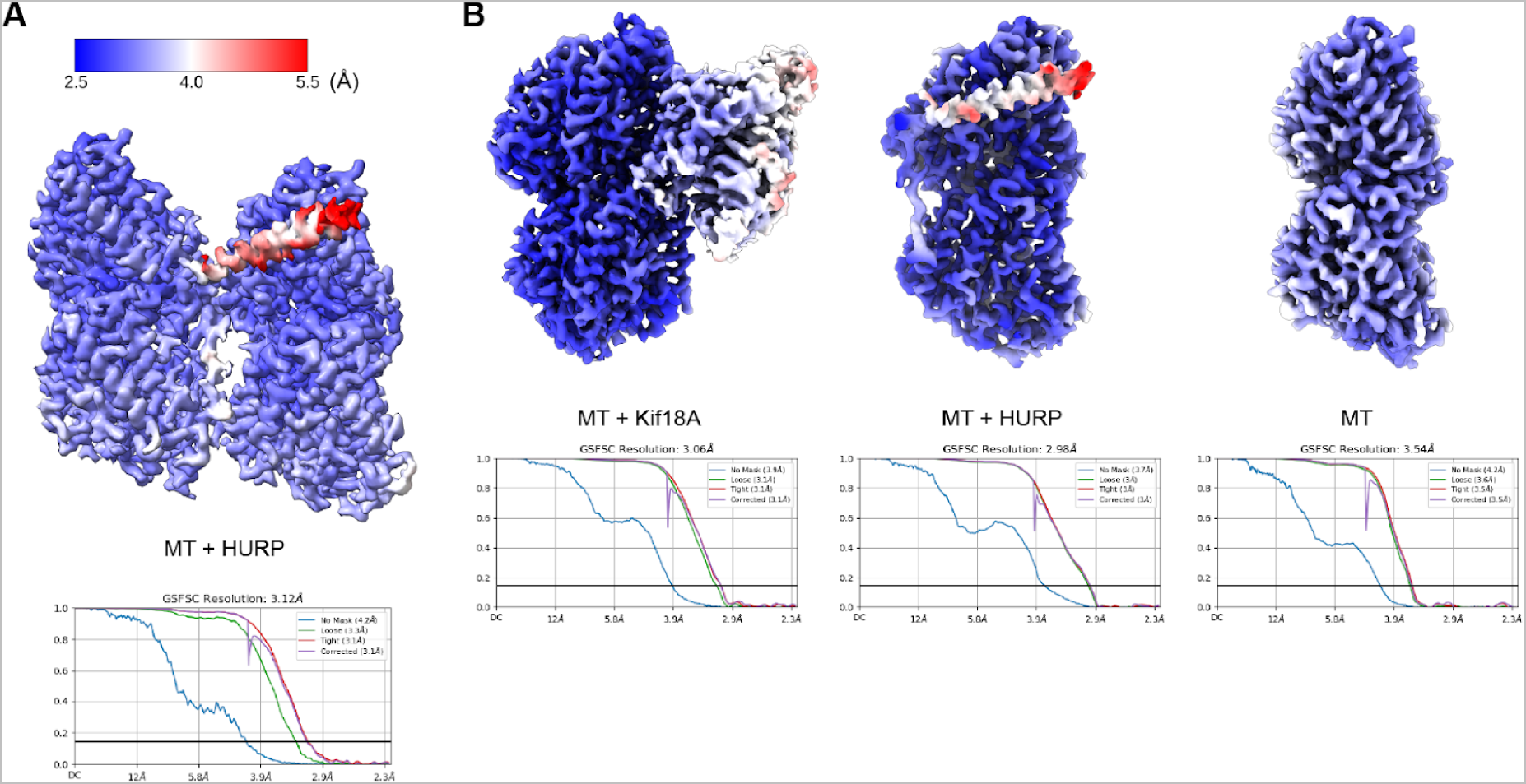
Local resolution maps and FSC curves. **A.** Local resolution map (top) and Fourier Shell Correlation (FSC) plots (bottom) for microtubule-bound HURP. **B.** Local resolution maps (top) and Fourier Shell Correlation (FSC) plots (bottom) for the three classes produced after 3D classification of the microtubule-HURP-Kif18A^1-373^ dataset.

**Supplementary Figure 6.**
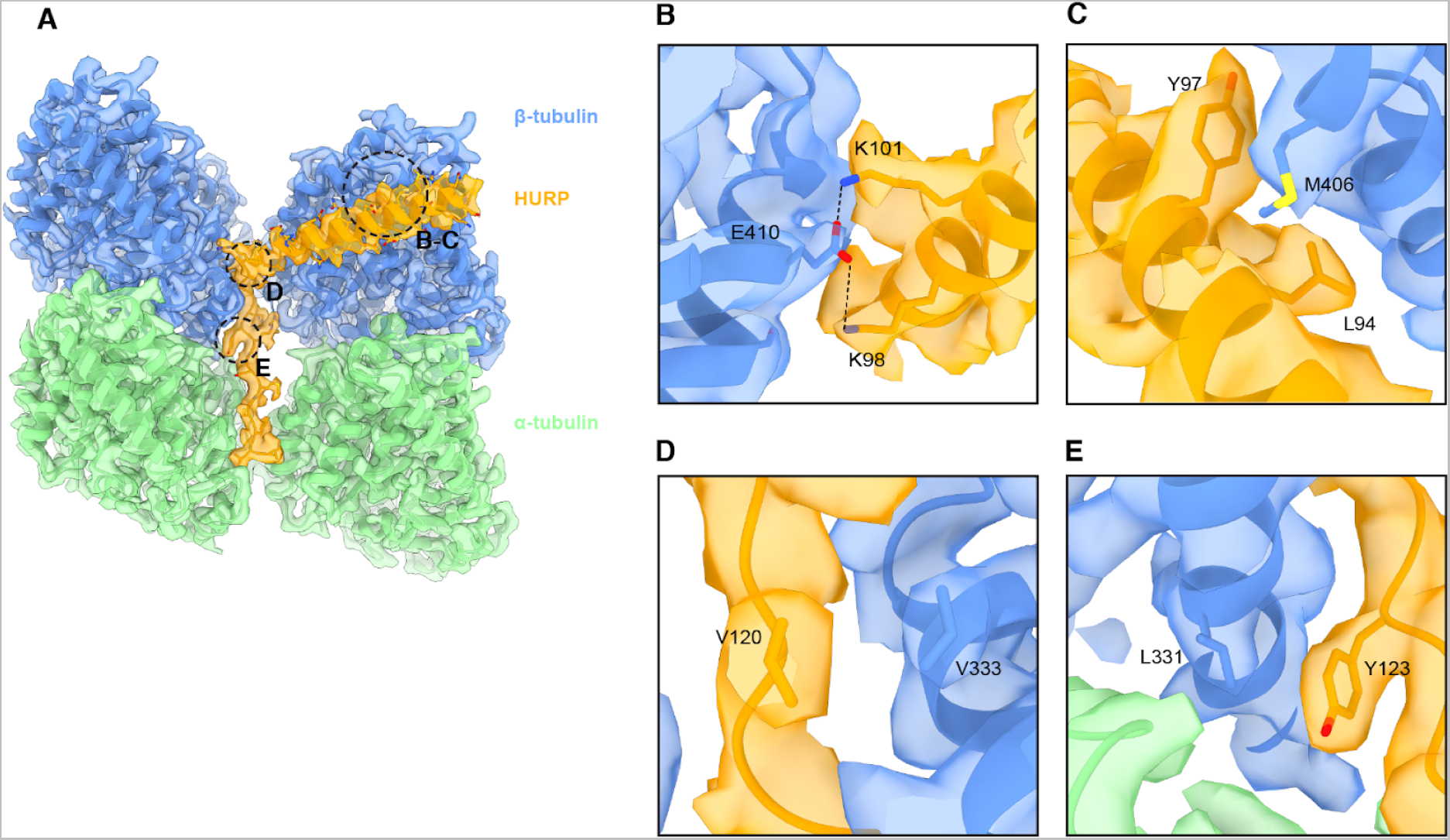
Detailed interactions between HURP and tubulin. **A.** Final microtubule-HURP cryo-EM map. A single HURP molecule is shown for clarity. The refined model is overlaid on the map. Dashed circles indicate the regions shown in panels B-E. **B-E.** Additional interactions between HURP and tubulin.

**Supplementary Figure 7.**
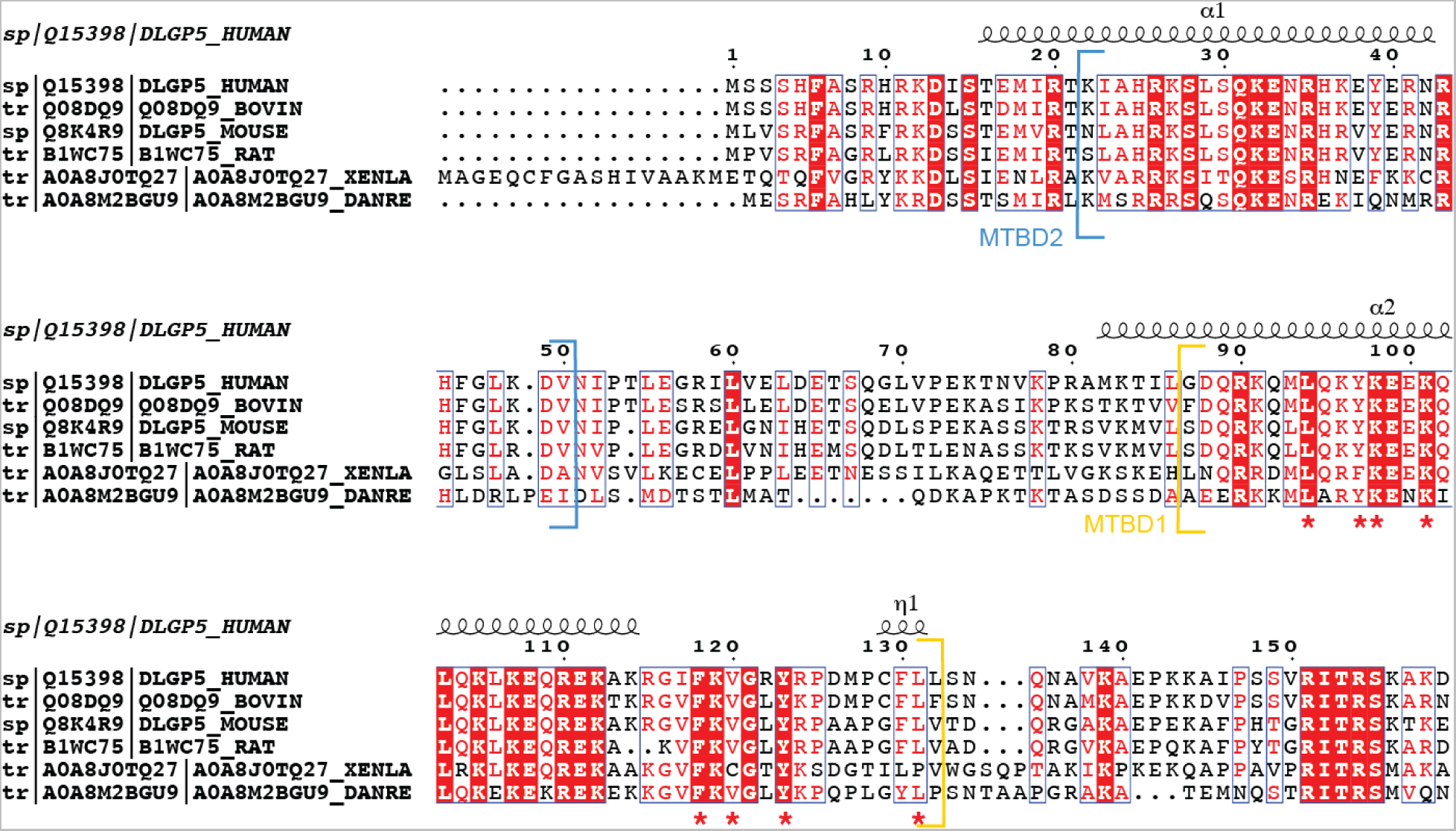
HURP residues that interact with tubulin are conserved across species. Sequence alignment and conservation analysis for HURP from different species. The human version was set as a reference and residues that are seen to interact with tubulin in our cryo-EM structure are marked with red asterisks. ESPript3^70^ was used to generate the figure, and secondary structure was annotated based on the AlphaFold^71^ prediction for human HURP (Uniprot Q15398). Regions corresponding to MTBD2 (blue) and the structurally resolved part of MTBD1 (yellow) are indicated.

**Supplementary Figure 8.**
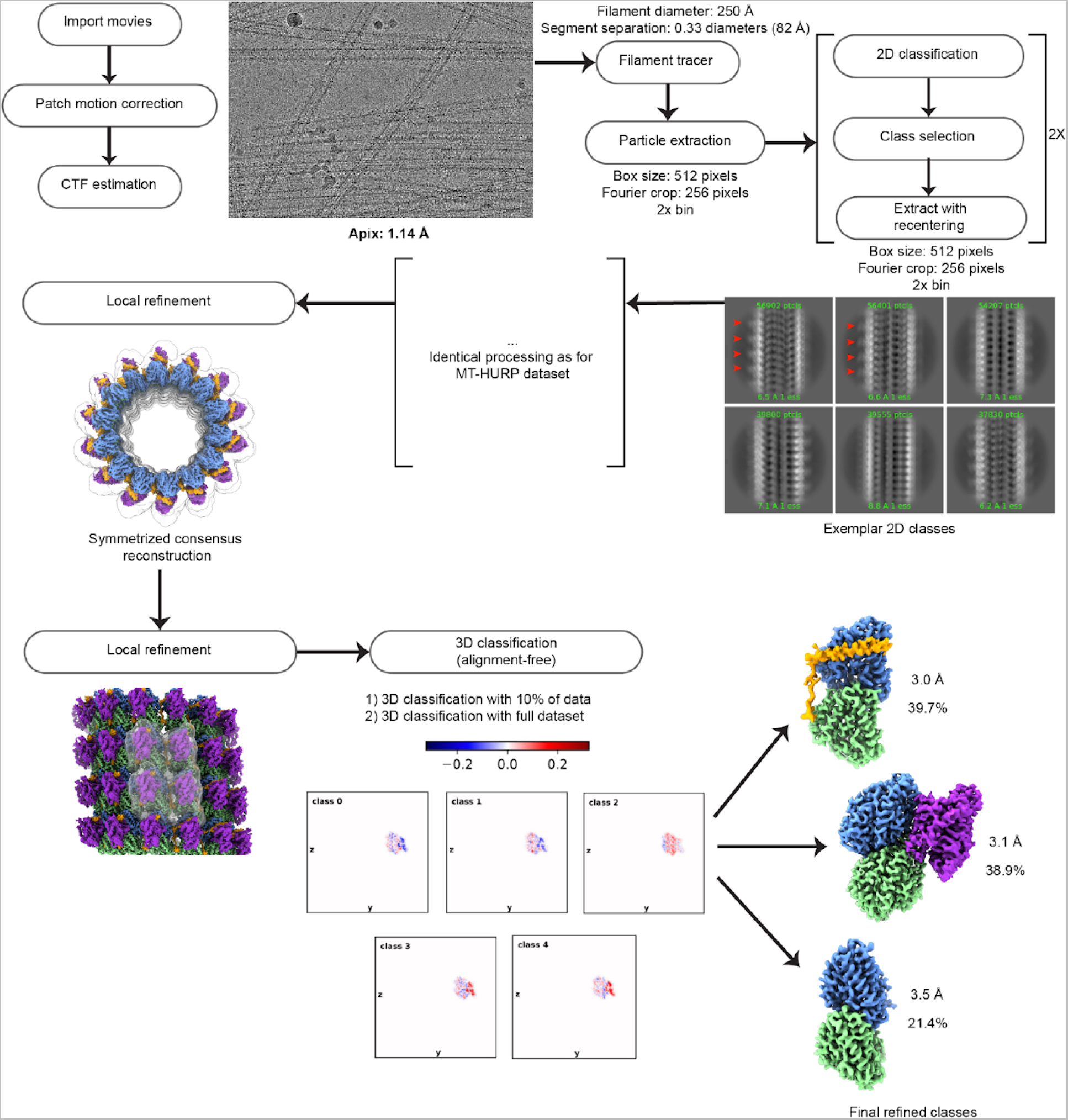
Cryo-EM data processing for the microtubule-HURP-Kif18A dataset. Processing pipeline applied to the microtubule+HURP+Kif18A^1-373^ dataset. Unless specified otherwise, all steps were performed in CryoSparc. Red arrowheads point to Kif18A density in the 2D class averages. In the 3D classification step, the blue-red key represents density variation in each class with respect to the input consensus reconstruction, with regions in blue and red containing significantly less or more density than the consensus, respectively. Masks are shown with transparent surfaces.

**Supplementary Figure 9.**
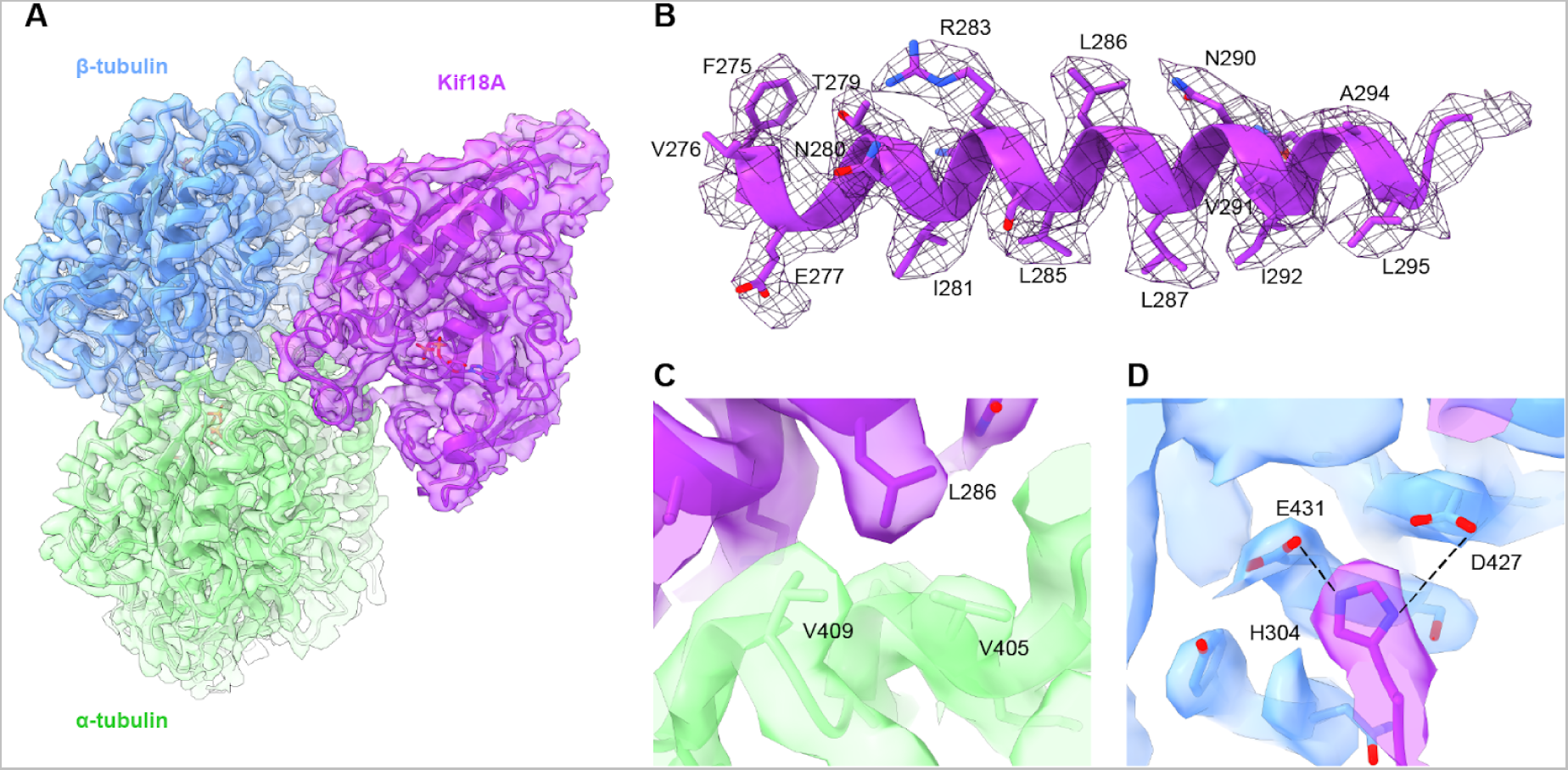
Map quality and detailed interactions for the Kif18A-tubulin class. A. Surface representation of the cryo-EM density corresponding to the microtubule+Kif18A class, with a refined tubulin+Kif18A model fitted inside the map. α-tubulin, 𝛽-tubulin and Kif18A are represented in green, blue and purple, respectively. **B.** Isolated density in mesh representation for Kif18A’s α4 helix, with the corresponding segment of the real-space refined atomic model of Kif18A. **C-D.** Selected detailed interactions between Kif18A and tubulin.

**Supplementary Figure 10.**
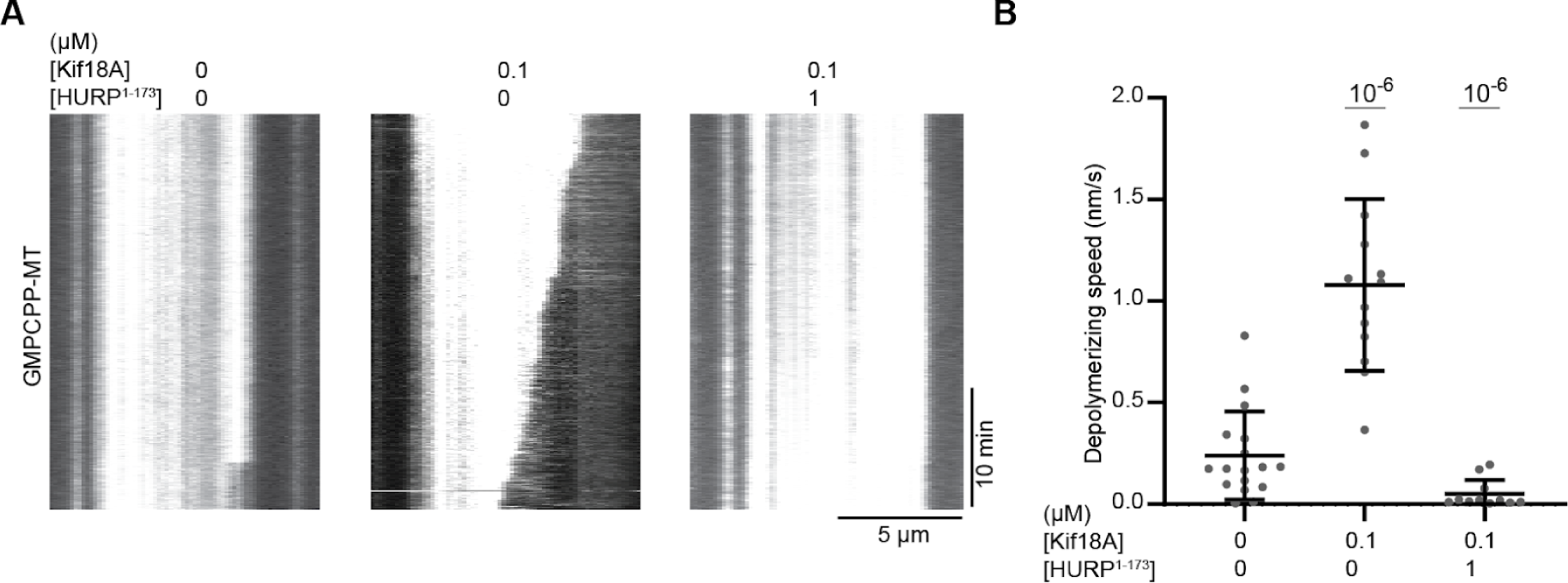
Kif18A can depolymerize GMPCPP microtubules. **A.** Representative kymographs of GMPCPP-microtubule depolymerization with 0.1 µM Kif18A or 0.1 µM Kif18A + 1 µM HURP^1-173^. **B**. Microtubule depolymerization speed with 0.1 µM Kif18A or 0.1 µM Kif18A + 1 µM HURP^1-173^. (From left to right, n = 17, 13 and 11 kymographs). The center line and whiskers represent the mean and S.D., respectively. P values are calculated from a two-tailed t test.

**Supplementary Figure 11.**
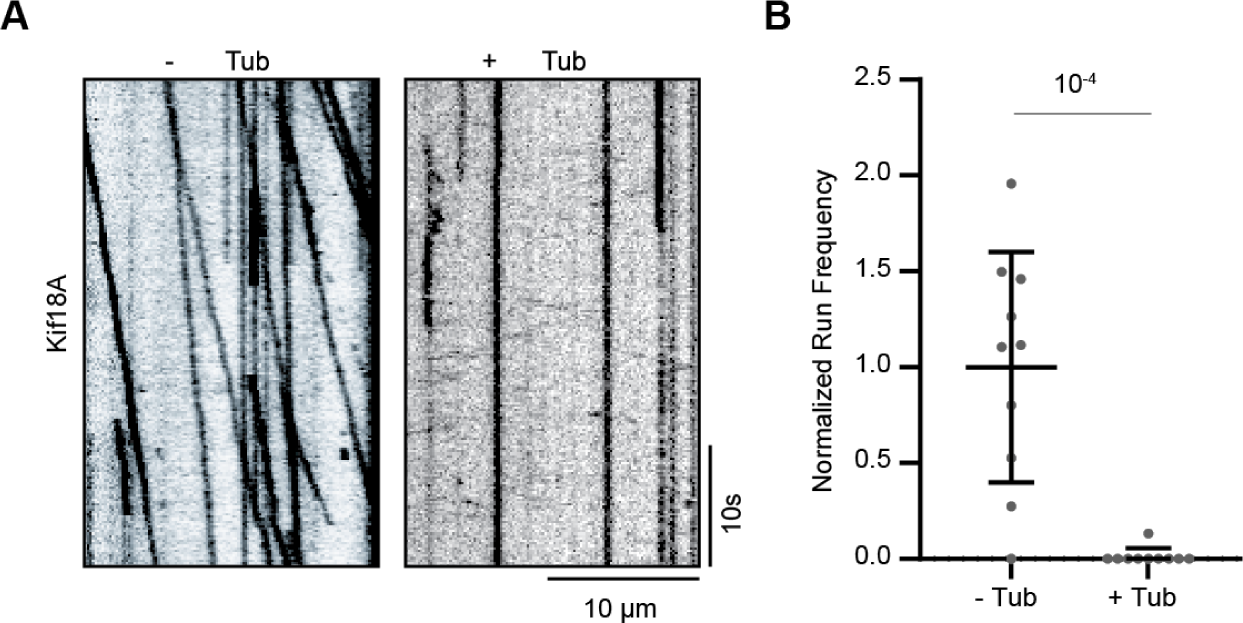
Kif18A motility is affected by the presence of free tubulin. **A.** Representative kymographs showing the motility of full-length Kif18A with or without 2 mg/mL free tubulin. **B**. Normalized run frequency of full-length Kif18A with or without 2 mg/mL free tubulin (n = 10 kymographs for each condition). The center line and whiskers represent the mean and S.D., respectively. P values are calculated from a two-tailed t test.

**Supplementary Figure 12.**
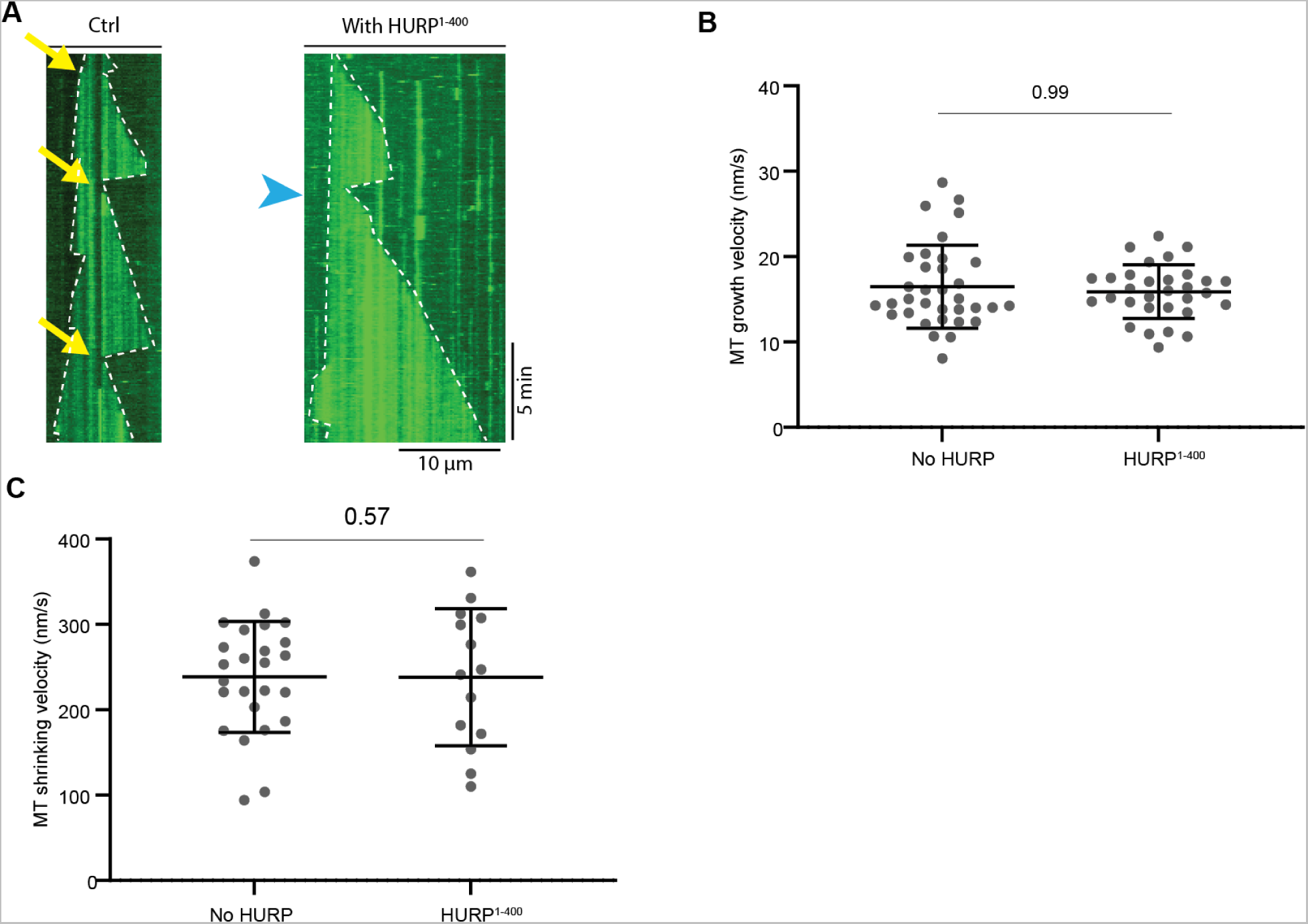
Effect of HURP^1-400^ on dynamic microtubules. **A.** Kymographs of dynamic microtubules with or without HURP^1-400^. Yellow arrows represent catastrophe events and blue arrowheads represent rescue events. **B.** Microtubule plus-end growth velocities with or without HURP^1-400^ (from left to right, n = 34, 31 microtubule growth periods). **C.** Microtubule plus-end shrinking velocities with or without HURP^1-400^ (from left to right, n = 25, 14 microtubule shrinking periods). In **B**-**C**, the center line and whiskers represent the mean and S.D., respectively. P values shown above the data points are calculated from a two-tailed t test.

**Supplementary Figure 13.**
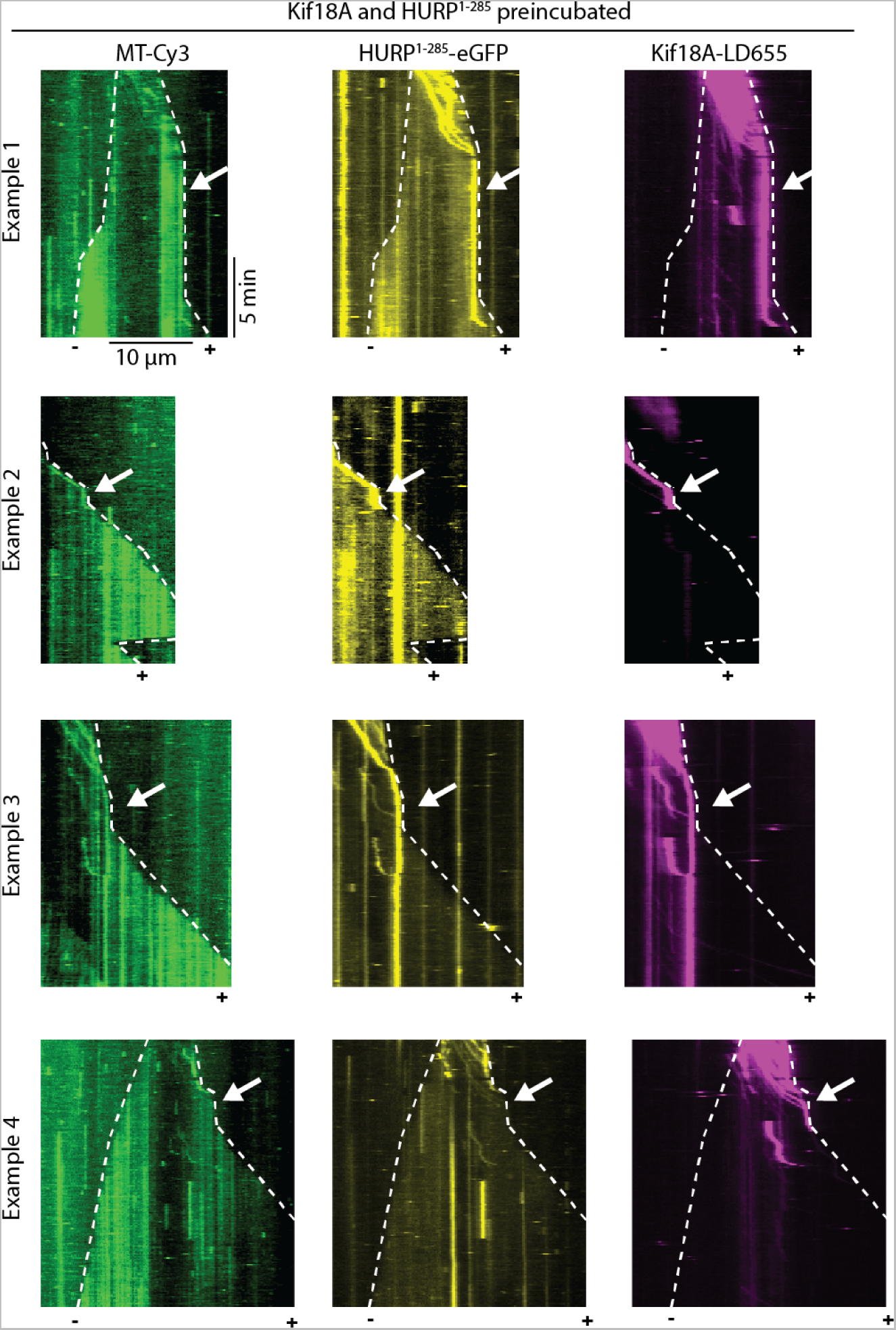
Kif18A and HURP jointly control microtubule length. Kymographs of Kif18A and HURP^1-285^ collectively maintaining a constant microtubule length (shown by white arrow). White dashed lines show the track of microtubule ends.

**Table S1.** Cryo-EM data collection parameters and model refinement statistics.

| Name | MT + HURP | MT + HURP + Kif18A (HURP class) | MT + HURP + Kif18A (Kif18A class) | MT + HURP + Kif18A (Tubulin class) |
| --- | --- | --- | --- | --- |
| EMDB ID | XXX | XXX | XXX | XXX |
| Microscope | Arctica | Arctica | Arctica | Arctica |
| Voltage (kV) | 200 | 200 | 200 | 200 |
| Camera | K3 | K3 | K3 | K3 |
| Defocus range ($\mu\text{m}$) | 0.8-2 | 0.8-2 | 0.8-2 | 0.8-2 |
| Automation software | SerialEM | SerialEM | SerialEM | SerialEM |
| Frames | 50 | 50 | 50 | 50 |
| Total dose (electrons/$\text{\AA}^2$) | 50 | 50 | 50 | 50 |
| Pixel size ($\text{\AA}/\text{pixel}$) | 1.14 | 1.14 | 1.14 | 1.14 |
| Number of micrographs | 796 | 2611 | 2611 | 2611 |
| Starting number of particles (pre-symmetry expansion) | 99,992 | 1,151,884 | 1,151,884 | 1,151,884 |
| Number of particles in final map (post-symmetry expansion) | 353,980 | 2,405,078 | 2,349,581 | 1,296,716 |
| Map sharpening B factor ($\text{\AA}$) | -60 | -100 | -60 | -80 |
| Map sharpening methods | CryoSparc | CryoSparc | CryoSparc | CryoSparc |
| Symmetry | C1 | C1 | C1 | C1 |
| Overall resolution ($\text{\AA}$) | 3.1 | 3.0 | 3.1 | 3.5 |
| Resolution range of map (min-75th percentile) ($\text{\AA}$) | 2.8-4.1 | 2.5-5.6 | 2.6-4.0 | 2.9-4.4 |

| Metric | Value |  |
| --- | --- | --- |
|  | MT + HURP | MT + Kif18A |
| Initial model used (PDB ID) | 6DPV | 5OCU |
| Refinement package | Phenix | Phenix |
| C-beta outliers (%) | 0 | 0 |
| Rotamer outliers (%) | 0.34 | 0.1 |
| All-atom Clash score | 9.17 | 8.76 |
| MolProbity score | 1.94 | 2.01 |
| Ramachandran plot<br>(outliers/favored) (%) | 0.64/93.00 | 0.44/90.39 |
| Ligand | 8 | 6 |
| Protein residues | 1724 | 1157 |
| R.m.s.d. of bond lengths (Å) | 0.003 | 0.004 |
| R.m.s.d. of bond angles (°) | 0.644 | 0.733 |
| CC (mask) | 0.84 | 0.81 |
| CC (volume) | 0.84 | 0.75 |

